# Breakthrough Percepts of *Familiar* Faces

**DOI:** 10.1101/2025.03.22.644718

**Authors:** Omid Hajilou, Howard Bowman

## Abstract

In Rapid Serial Visual Presentation (RSVP), the vast majority of stimuli are not consciously perceived, but the salient ones breakthrough into awareness and can be reported. In addition, these breakthrough events are observable with EEG, since they generate a P3 or other distinguishing components. The *Fringe-P3* method is based upon these characteristics. Concealed knowledge studies have successfully employed this *Fringe-P3* method using own-name and own email-address, with the method being shown to be less vulnerable to counter-measures than other approaches. It has also been shown that famous faces presented in RSVP differentially break into awareness and generate a distinct evoked response component.

In this paper, we further enhance the applicability of the *Fringe-P3* concealed knowledge test by demonstrating the effectiveness of the method on personally-familiar faces. While salient, such stimuli do not have the exquisite salience of famous faces, being a better match to the level of salience that might be found in forensic applications. Our findings suggest that the *Fringe-P3* method could be used to detect intrinsic salience of familiar faces, even when there was no task associated with these faces. We investigated the sensitivity of the ERP-based RSVP paradigm to infer recognition of familiar faces, and performed statistical inference in the Time and Frequency domains, to differentiate between known and unknown faces, at group and participant levels.

## INTRODUCTION

The classic oddball P3 experiments have been successfully used in an investigative context as Concealed Information Tests (CIT) (Johnson & Rosenfeld, 1992) (Rosenfeld, Soskins, Bosch, & Ryan, 2004) and (Labkovsky & Rosenfeld, 2012). Additionally, CITs have successfully used face stimuli in these P3-oddball paradigms (Meijer, Smulders, & Wolf, 2009) and (Lefebvre, Marchand, Smith, & Connolly, 2007). However, these P3-oddball paradigms have exhibited a vulnerability to countermeasures (Rosenfeld, Soskins, Bosch, & Ryan, 2004). Recent CIT work with Rapid Serial Visual Presentation (RSVP) (Bowman, et al., 2013); (Bowman, Filetti, Alsufyani, Janssen, & Su, 2014) has been argued to be less vulnerable to countermeasures. In particular, salient stimuli that are presented in RSVP streams (e.g. a familiar name) will breakthrough into awareness, whilst non-salient stimuli (e.g. an unknown name) will not enter awareness making it more difficult for participants to use countermeasures. For example, where stimuli are presented at a slow rate, suspects could confound the tests by using countermeasures (Meixner, Haynes, Winograd, Brown, & Rosenfeld, 2009); (Lukács, et al., 2016). A key reason for the possibility of countermeasures is the availability of sufficient time between each stimulus, to consciously determine that there is a repeating non-critical irrelevant item^1^. However, (Bowman, Filetti, Alsufyani, Janssen, & Su, 2014) observed that if the stimuli were presented at a rapid rate in RSVP, the irrelevant stimulus was not sufficiently perceived to be noticed as repeating, preventing it being used in a countermeasure. This difficulty for participants to detect non-salient repeating stimuli has been confirmed in a range of empirical work (Avilés, Bowman, & Wyble, 2020); (Bowman & Avilés, 2021); (Bowman & Avilés, 2022).

These RSVP findings were implemented in the *Fringe-P3* method, a brain computer interface, which emerged from theoretical work on the attentional blink (Bowman & Wyble, 2007); (Bowman H., Wyble, Chennu, & Craston, 2008); (Craston, Wyble, Chennu, & Bowman, 2009). The method involves presenting an RSVP stream and determining the stimuli in the stream that the participant finds salient by detecting a “breakthrough” event, i.e. the participant’s perceptual system detecting a salient stimulus and accordingly its representation being consciously perceived. This perceptual event manifests as a P3 or other distinguishing EEG component. The method has been used as a P3-speller, a classic brain computer interface (Chennu, Alsufyani, Filetti, Owen, & Bowman, 2013) and as a CIT.

Having provided evidence for the effectiveness of the Fringe-P3 CIT based upon own-name (Bowman, et al., 2013); (Bowman, Filetti, Alsufyani, Janssen, & Su, 2014), own email-address (Harris, et al., 2021), famous name (Alsufyani, Harris, Zoumpoulaki, Filetti, & Bowman, 2021), future experiments were required to explore the suitability of image-based stimuli, in ERP-based RSVP paradigms; for example, do familiar faces differentially break into conscious awareness, on an individual basis, and can we detect the breakthrough events with EEG? To answer this question, we began by studying the effects of presenting famous faces (i.e. Probe stimuli) and unknown faces (i.e. Irrelevant stimuli), using the RSVP technique.

Accordingly, we are able to successfully establish that famous/celebrity faces breakthrough into conscious awareness, using an RSVP subliminal search paradigm, and that our statistical tests can differentiate between the Probe (famous) and Irrelevant (unknown) faces, at group and participant levels (Alsufyani, et al., 2019). The objective of the current paper is to demonstrate that we can substitute the highly evocative faces of famous celebrities with familiar faces that are personally known to the participants. While salient, such stimuli do not have the exquisite salience of famous faces, and are thus a better match to the level of salience that might be found in forensic applications. Additionally, we assessed whether the *Fringe-P3* method could be used to detect intrinsic salience of familiar faces, even when there was no task associated with these faces.

ERP studies into face perception and recognition have reliably reported neural activity, with specific components that indicate sub-conscious and conscious processing of faces. Numerous studies have reported a face-specific N170 component – a negative deflection, elicited in the ERP within 140 and 200ms post-stimulus.

Whilst the N170 component does not appear to be affected by the difference between famous and unfamiliar faces (Bentin & Deouell, 2000), an enhanced negativity called the N400 component (also referred to as N400face, or N400f) appears to be associated with the subsequent activation of the episodic memory of the face (Bentin & Deouell, 2000); (Touryan, 2011); (Eimer, 2000). Amongst others, (Eimer, 2000) observed a subsequent/late positivity for famous faces, referred to as the P600 component (or P600f), which has also been compared with the P3 component (Sutton, Braren, Zubin, & John, 1965). This P3/P600f neural response may reflect explicit recognition of a particular individual.

In (Alsufyani, et al., 2019), we used the N400f and the P600f for our time domain analyses of the brain response to famous faces presented in RSVP both at the group and per-individual level. However, our most effective per-individual statistical test in (Alsufyani, et al., 2019) was in the frequency domain, where we performed a permutation test over power and inter-trial coherence features. These time and frequency domain analyses will be reused in this paper exactly as used in (Alsufyani, et al., 2019) to differentiate between the Probe (familiar: lecturer at participant’s institution, University of Kent) and Irrelevant (unknown: lecturer at different institution, Christchurch University). This gives us strong prior precedents for the analyses we perform here, providing robust protection against type 1 errors.

The aim of this work is, then, to provide a proof of concept, which could be further refined and developed into a scientifically robust framework, in the pursuit of a means by which a suspect’s familiarity with compatriots can be demonstrated using EEG.

## METHODS

### Participants

Fourteen participants were tested, who were all students at the University of Kent, free from neurological disorders, and with normal, or corrected-to-normal vision. None were excluded. Out of 14 participants, 12 were male (86%) and 2 female (14%). The ages of the participants ranged from 22 to 37 (*M* = 27.5 years, *SD* = 3.94); 12 of them were right-handed (86%), and 2 were left-handed (14%). Because this experiment’s Probe stimuli consisted of University of Kent’s School of Computing lecturer faces, we needed participants who had a close working relationship with these lecturers. Therefore, we asked the lecturers to covertly nominate PhD students, so that we could be assured of a long-term relationship/familiarity between participants and their lecturers’ faces. Also, participants were chosen on the basis of never having been included in a similar EEG/RSVP experiment, and at the end of each experiment, participants were instructed to avoid discussing the experiment with their colleagues, in order to avoid any priming of future participants. The duration of each experiment was (approx.) 1 hour and 45 minutes, and each participant was paid £10 (ten pounds) for their time. The Sciences Research Ethics Advisory Group, at the University of Kent, approved the study.

### Stimuli

The instructions, stimuli and questions were presented on a 20-inch LCD monitor, with a refresh rate of 60Hz, and a resolution of 1600×1200 pixels. The screen was placed at a comfortable position for each participant, at a distance of approximately 60 to 80cm. All stimuli were scaled to 280×320 pixels, and presented in the centre of the screen, using the Rapid Serial Visual Presentation (RSVP) method.

The stimuli were split into two groups: a) **Distractor** images (i.e. unknown faces) and b) **Critical** images, as described below:

**a) Distractor** images (i.e. the first group) were photographs of unfamiliar faces, which were obtained from an open-source, online database of faces (Minear & Park, 2004) from the University of Texas at Dallas – all of these faces were frontal views.
**b) Critical** images (i.e. the second group) consisted of the following three categories:

**i) Target** face, which was a single image, chosen by us from the Distractors database. The Target was task-relevant (i.e. participants were instructed to look out for, and respond to this image).
**ii) Irrelevant** (a.k.a. *unknown*) faces consisted of unknown faces of Lecturers from another University.
**iii) Probe** (a.k.a. *familiar*) faces consisted of several images of familiar faces, in the form of our University’s lecturer faces.

Care was taken to ensure that all the images used in the experiment would conform to our compatibility criteria. Accordingly, images with incongruous features (e.g. angry or smiley faces), which could breakthrough into conscious awareness due to their dissonant features, were avoided. As a result, the large database of Distractor faces (with over 1000 images) was reduced to 573 ‘neutral’ images, without significant facial expressions or features. The remaining 573 Distractor images were available as fillers for RSVP streams, albeit, one was selected as the Target, for the experiment.

The Probes (i.e. familiar Lecturer faces from the University of Kent’s School of Computing), as well as, the Irrelevants (i.e. unknown Lecturer faces from Christ Church University) were taken using the same SLR camera (Canon PowerShot G11). All lecturers consented to their image being used in our EEG experiments; these photos were taken from the same position/distance, under similar lighting conditions, and with neutral poses.

All images of lectures were manually edited, in order to remove any non-conforming distinguishing features, and to obtain similar brightness and contrast. First, each image was centred by aligning the eye-line to the same horizontal position, and then resized, to occupy the same space/size as the Distractor images. Next, the background of each lecturer face was removed (i.e. borders were carefully highlighted/selected, using the photo-editing tool GIMP, before being cropped out), and then the background colour was changed to light-grey (Hex colour: #e7e7e7). Next, the contrast of the images were reduced, wherever necessary, and all Probe images were resized to 280×320 pixels and converted to monochrome.

After the above exercise to select-and-edit our Probe images, we decided to further reduce the Distractor images to 524 faces, in order to approximately match the age range of the Probe faces (i.e. by excluding Distractor images of very young and very old individuals). As there were 524 possible Distractors, which could be used as fillers for RSVP streams, the probability that one would be randomly selected for each stream was 0.032, and equal for all of them.

### Probe/Irrelevant comparison

As explained above, Probe and Irrelevant critical stimuli were lecturer faces that were photographed by us. Having photographed a large portion of the lecturers in the University of Kent’s School of Computing (23 images in total), we were able to assign three Lecturers (as Probes) to each participant, knowing that they were familiar with one another (as confirmed by the Lecturers and/or the participant’s colleagues). Additionally, each participant was randomly assigned three unknown lecturers (as Irrelevants) from a different University (i.e. Canterbury Christ Church University).

### Design

All stimuli were presented using the *Psychophysics* toolbox version 3, running under MATLAB 2012a. All RSVP stream items were presented in the same location (centre of the screen), at an SOA of 133ms (see Figure 1), and without an Inter-stimulus-Interval (ISI). Each RSVP stream contained 18 faces, 17 of which were Distractors (i.e. unknown fillers), and only one was a Critical item (i.e. either a Probe, Irrelevant or Target). To ensure that neither the Probes nor the Irrelevants were task-relevant, we instructed all participants to look for a single Target (i.e. an unknown face that participants were trained to detect), throughout the experiment. This task-relevant Target was repeated as many times as each of the other two Critical stimuli (i.e. the Target, Probe and Irrelevant conditions were repeated equal number of times) and (in a statistical sense) in the same position in streams.

**Figure 1.**
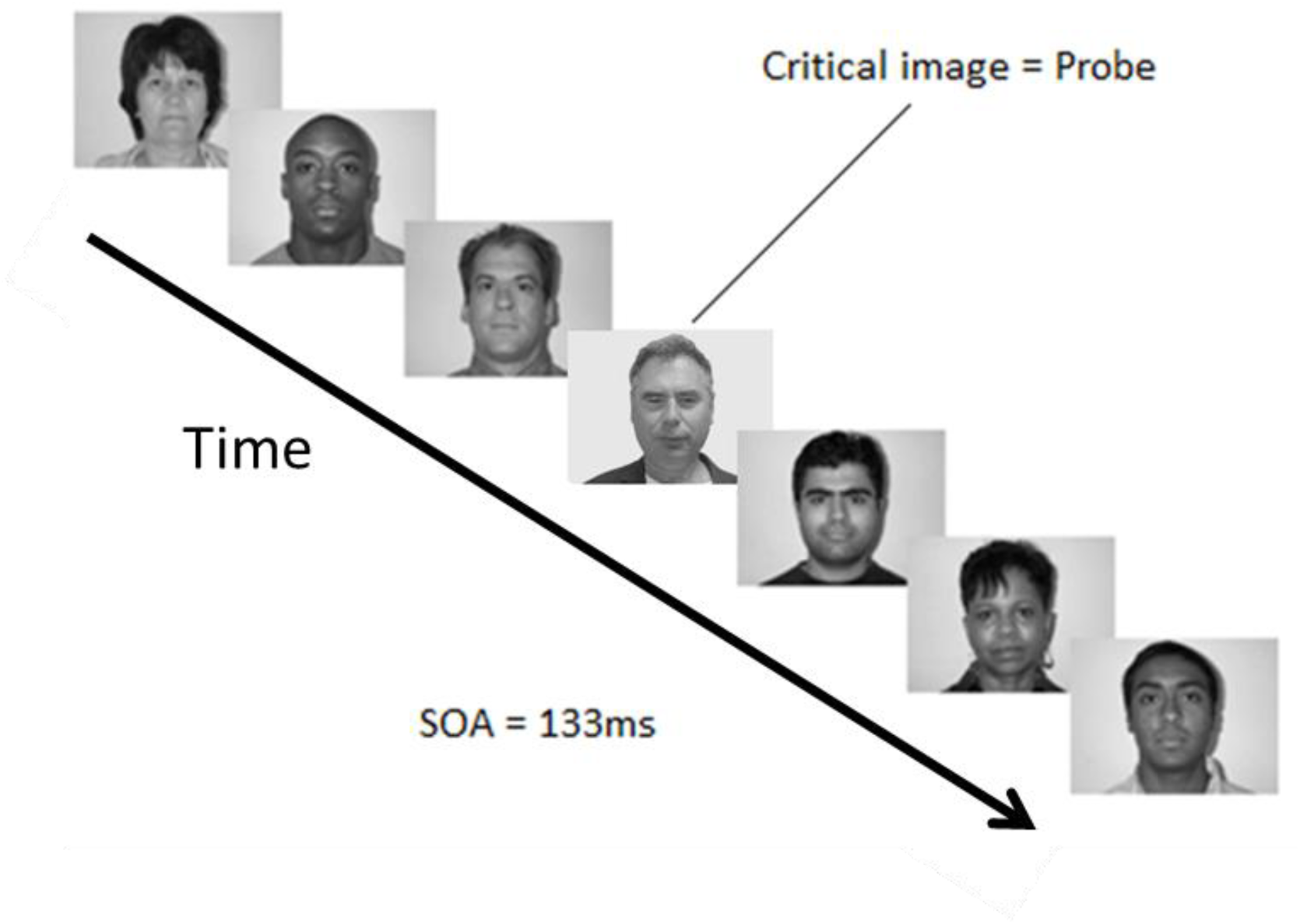
RSVP stream, showing 7 of the 18 faces that could be presented in a trial, where each trial consists of 17 Distractors (i.e. unfamiliar faces), and one Critical image (i.e. a Probe, Irrelevant or Target). In the above example from the experiment, a Lecturer (Howard Bowman) is the Probe that is presented as the Critical image.

We implemented a change to the design of the famous faces experiment (Alsufyani, et al., 2019): to improve the signal-to-noise ratio of unique Probe/Irrelevant Critical stimuli, we reduced the number of Probes/Irrelevants from five (celebrities) to three (lecturers), whilst maintaining the total number of trials for each participant (i.e. 225 trials in total, so that the number of trials per Critical condition would also remain the same between the two experiments). Therefore, in this experiment, each Probe was repeated 25 times, resulting in 75 Probe-trials (i.e. 25 times for each of the 3 Probes), and each Irrelevant was also repeated 25 times, resulting in 75 Irrelevant-trials (i.e. 25 times for each of the 3 Irrelevants). The single Target was also repeated 75 times, to equal the number of repetitions of the other two Critical Stimuli. The resultant 225 RSVP trials were divided into 3 blocks, each block comprising 75 trials (i.e. 25 Probe trials, 25 Irrelevant trials, and 25 Target trials), and the order of the three Critical stimuli were randomised within the blocks. However, each block’s Probe and Irrelevant Critical stimuli were paired, so that the same known lecturer (Probe) face and unknown lecturer (Irrelevant) face were presented within the same block – this will enable us to make direct comparisons between these paired-conditions.

Participants were told to keep their eyes fixed at the centre of the screen during the presentation of the RSVP stream (lasting approx. 2.5s), and to avoid movement or blinking. Also, they were informed that the Target image will appear pseudo-randomly, so they should not expect it in every trial, however, participants were naïve to the presence of familiar lecturer faces (i.e. Probes).

In this experiment, out of a total of 75 trials for each Critical condition, the number of trials that remained after artefact rejection, per condition, ranged between 59 and 75, and none of the participants were excluded from the analysis due to removal of artefact trials (e.g. excessive eye blinks):

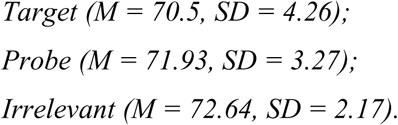

The position of the single Critical item within each RSVP trial, which contained 17 Distractors as fillers, was selected pseudo-randomly by the application, so that it had equal probability of appearing anywhere in the 5^th^ to 9^th^ position of the stream of images. The starting boundary (i.e. the first 4 items of the stream) was avoided because of stream onset transients. Similarly, the ending boundary (i.e. the last 9 items of the stream) was also avoided because of offset transients and anticipation of the end of stream item-and-question.

In addition to the 18 images, each RSVP stream contained a starting and finishing item, which would improve the participants’ focus, from the beginning to the end of each stream (i.e. to know when the stream is about to start, and when it will end). A starting item “+ + + + + + +” was presented for 800ms, to position the participant’s focus on the presentation area of the screen. After the last image of the RSVP stream (i.e. the 18^th^ face), a random finishing item (a.k.a. attention-checker image), which could either be “ ” or “= = = = = = =”, was presented for 133ms; this end-of-stream image required the participant to remain attentive until the end of the stream. Using a standalone keypad, which was positioned under the participant’s preferred left or right hand, the participant was asked to report the attention-checker item, using ‘1’ and ‘2’ keys (1 signifying “ ” and 2 signifying “= = = = = = =”).

As soon as the participant responded to the attention-checker question, a task-relevant question was shown, in order to determine the detection (or not) of the Target image, using the question: “Did you see the *Target* image within the stream?”. In response, the participant could select the keys ‘4’ (for “Yes”) or ‘5’ (for “No”). The above two questions were repeated at the end of each RSVP stream. Recognition-questions were also included at the end of the experiment. These were included at experiment end in order to mitigate the participants’ inference that the Probe (which would have been in the recognition test) could also be task-relevant (i.e. in addition to the Target), which could have occurred if recognition was tested during the course of the experiment.

Participants were given one (or more) training session(s). Each training session consisted of 20 RSVP trials, to ensure that the participant was comfortable observing the rapid presentation of images, and to make sure that they could identify the Target. During the training session(s), the RSVP streams did not contain any images that were assigned to the Probe or Irrelevant category of critical stimuli in the experiment proper.

### Analysis

#### Data acquisition

The experiment was recorded using a BioSemi ActiveTwo system (*BioSemi, Amsterdam, The Netherlands; see* www.biosemi.com). The Electroencephalographic (EEG) data was filtered at recording, with a low-pass of 100 Hz, and digitised at 2,048 Hz, for offline analysis, and the impedance was kept below 10 kΩ. In accordance with the standard 10-20 system (Jasper, 1958), the following 8 scalp electrodes were used in the first two experiments: Fz, Cz, Pz, P3, P4, Oz, A1 and A2. During recording, data was referenced to a ground formed from a Common Mode Sense (CMS) active electrode, and Driven Right Leg (DRL) passive electrode. These two electrodes form a feedback loop, which drives the average potential of the participant, as close as possible, to the Analogue to Digital Converter (ADC). Also, Electrooculograms (EOG) generated from eye blinks and eye movements, were recorded from the participant’s left and right eyes, using two bipolar Horizontal EOG (HEOG) and Vertical EOG (VEOG) electrodes.

#### EEG data

For all three experiments, the recorded data, which was analysed using EEGLAB (version 12.02.4b), under MATLAB 2012a (Delorme & Makeig, 2004), was resampled at 512 Hz. Before analysis, we filtered the data with low-pass, and high-pass, filters of 45 Hz to 0.5 Hz, respectively. Furthermore, in order to remove the steady state Visually Evoked Potential (ssVEP) oscillations (Wang & Jung, 2010), notch filters were applied between 7 Hz and 9 Hz. Then, we re-referenced the data off-line to the average of the combined mastoids (i.e. A1 and A2 electrodes), and generated ERPs by separately averaging all trials for each Critical condition (i.e. Target, Probe, and Irrelevant). Each EEG trial was generated by epoching the data, using a -200ms to 1200ms stimulus-locked window (i.e. -200ms before the appearance of the Critical stimulus, and 1200ms after the occurrence of the Critical stimulus), and all ERPs were time-locked to the onset of the Critical item (i.e. time-point zero marks the appearance of the Critical face). Although baseline correction could be applied at trial level (i.e. mean of -100ms to 0ms window subtracted from each trail), the new standard for applying the detrending technique (see Application of Detrending) required us to baseline correct each trial after the adjustment for any errant drift.

Eye blinks and muscle movements were detected, by marking any activity below - 100µV or above +100µV in the EOG channels (reflecting eye blinks and horizontal/vertical movements). Furthermore, we rejected any trials containing electrical activity below -50µV or above +50µV, in a time window from -200ms to 1200ms (reflecting other physiological and environmental artefacts), with respect to the Critical stimulus Onset. Finally, we performed manual inspection of the resulting ERP data, to verify that the rejected trials were accurately detected.

#### Time Domain (ERP) Analysis

Each experiment’s trails can be split into three categories: those with the *Target* critical stimulus, ones with *Probe* critical stimulus and the ones with *Irrelevant* critical stimulus (which was paired with the Probe). Our analyses were performed at the ERP-level, in order to compare the responses to familiar (Probe) and unfamiliar (Irrelevant) faces. As previously discussed, we focused on two face related ERP components: 1) an enhanced negativity called **N400** – also referred to as **N400f** (Curan & Hancock, 2007); and 2) a late positivity called **P3** – also referred to as **P600f** (Trenner, Jentzsch, & Sommer, 2004).

The time window associated with the above two ERP components should be identified based on a contrast independent of (strictly parametrically contrast orthogonal to) the contrast of interest (Brooks, Zoumpoulaki, & Bowman, 2017); (Bowman, Brooks, Hajilou, Zoumpoulaki, & Litvak, 2020). Thus, for pre-participant analysis, we selected the time window using the participant’s aggregated ERP, generated from all trials (hereafter called **aERPt**), within the combined Probe-and-Irrelevant conditions; this is also called the aggregated ERP of trials (Brooks, Zoumpoulaki, & Bowman, 2017); (Bowman, Brooks, Hajilou, Zoumpoulaki, & Litvak, 2020). Similarly, for group-level analysis, we selected the time window using an Aggregated Grand Average from all Trials (**AGAT**), belonging to all participants’ combined Probe-and-Irrelevant conditions (Brooks, Zoumpoulaki, & Bowman, 2017); (Bowman, Brooks, Hajilou, Zoumpoulaki, & Litvak, 2020). The following gives more details of the aERPt and AGAT.

#### Per-participant (aERPt) window placement and inference

For each participant, their aggregated ERP of all trials from both conditions, Probe and Irrelevant, was generated. This aERPt was then used to identify the time window of the two components of interest (i.e. N400f and P600f). For the N400f component, the entire ERP window (i.e. from 0ms to 1200ms) was searched to find the *minimal* 100ms interval average. Similarly, for the P600f component, the entire ERP window (i.e. from 0ms to 1200ms) was searched to find the *maximal* 100ms interval average. (We searched in the entire window, in order to give as much flexibility as possible for the latency of participant’s brain responses to vary and also, to bring as few (potentially biasing) constraints as possible to our analysis.) The start and the end of this minimal/maximal 100ms Region of Interest (ROI) defined the face related N400f and P600f time features/components.

After identifying the time windows for each component (e.g. the ROI for N400f, and the ROI for P600f), the mean amplitude was calculated separately to each condition within these time windows – in other words, for each component, one mean amplitude value for the Probe, and another for the Irrelevant, was calculated using the same time window that was independently found when Probe and Irrelevant trials were combined. The ‘True Observed’ difference of each component (i.e. N400f and P600f) in their respective ROI, is the difference between this measure for each condition:

#### True Observed difference = mean of Probe (minus) mean of Irrelevant

Having found the True Observed difference for N400f and P600f, we performed statistical inference, to determine whether the evoked response by the Probe (the familiar face) was significantly different from that evoked by the Irrelevant (the unknown face). Per-individual analysis was based on analysing each experimental participant separately to determine whether there was a significant difference for that participant alone. In this analysis, a randomisation (i.e. Monte Carlo Permutation) test was used to determine a *p*-value for each participant (see section “Randomisation Test”, below). A null hypothesis distribution for each participant was generated in order to calculate the individual’s *p*-value; the *p*-value estimates the probability that the observed pattern could have arisen if the null hypothesis was true, i.e. the participant was not familiar with Probe face.

#### Group-level (AGAT) window placement and inference

Time window placement for the group starts with the collation of all the trials for all participants, in both conditions (Probe and Irrelevant). The resultant Aggregated Grand Average of Trials (AGAT) was then used to identify the time window in the entire ERP segment of the two components of interest (i.e. N400f and P600f). For the N400f component, the *minimal* 100ms interval average was found, and for the P600f component, the *maximal* 100ms interval average was found. The start and the end of this minimal/maximal 100ms Region of Interest (ROI) defined the group-level face related N400f and P600f features/components. Finally, the mean amplitude measure was calculated separately for each condition, within the identified time window, and the ‘True Observed’ difference of each component (i.e. N400f and P600f) was obtained by finding the difference between each condition (similarly to in section “Per-participant (aERPt) window placement and inference”).

Having found the True Observed difference for N400f and P600f, we performed statistical inference, to determine whether the evoked response for the Probe (the familiar face) was significantly different from that evoked for the Irrelevant (the unknown face). A paired *t*-test of the mean amplitudes of Probe N400f/P600f and Irrelevant N400f/P600f was then run across participants.

#### Randomisation test

At per-individual-level, to determine whether the difference between two conditions (Probe and Irrelevant) is significant, we applied a randomisation (i.e. Monte Carlo Permutation) test. This was done separately for N400f and P600f components, for each participant. Before applying the test at the per-participant-level, the smallest number of trials between the Probe and Irrelevant conditions was determined and denoted by ‘*m*’ (they could contain different numbers of trials due to artefact rejection). Thus, only ‘*m*’ trials were selected (at random without replacement) from the Probe condition, and ‘*m*’ trials from the Irrelevant condition. Notably, if a direct comparison was to be made between individual blocks (as relating to a single familiar face), we made sure that the pairing of the Probe and Irrelevant conditions were maintained. Next, we calculated the difference between the mean amplitude values of Probe and Irrelevant ERPs (*Probe [minus] Irrelevant*), in order to obtain a mean amplitude difference measure. This mean amplitude difference became the **True Observed Value**.

The randomisation test was applied by populating a matrix of size (*2.m × number of time points*) with *2.m* selected trials; row position was randomised in the matrix. Under the null hypothesis, the Irrelevant and Probe trials are samples from the same distribution (i.e. the null distribution), and would thus be exchangeable. This justifies the randomisation of position in the matrix. Next, a pair of datasets were generated: the first, the surrogate Probe, was generated from the first half of the matrix rows, and the second, the surrogate Irrelevant, was generated from the remaining half. The desired analysis (i.e. mean amplitude measure) was then applied to each of the two randomised data sets, and the mean amplitude value was calculated (referred to as *Surrogate Values*), in the same way that the True Observed Value was calculated.

The above randomisation procedure was repeated 1,000 times. In each iteration, a new mean amplitude difference was obtained, resulting in 1,000 *Surrogate Values*. The *p*-value was then calculated as the proportion of randomised results that were greater than the true observed value. Finally, if this *p*-value is smaller than a critical alpha-level (0.05), then the data in the two experimental conditions are significantly different, and the null Hypothesis (H0) can be rejected. Note that in each resampling, the randomised mean amplitude difference (i.e. surrogate value) was measured for both the N400f and the P600f components (i.e. these values were calculated from the same random sample, rather than being calculated from two separate randomisations).

#### Frequency Domain Analysis

Previous studies into familiar faces (Bentin & Deouell, 2000) have reported ongoing oscillations in ERPs, from about 100ms to 500ms post stimulus onset, over parietal and occipital sites. In our famous faces experiment (Alsufyani, et al., 2019), we also observed multi-cycle oscillations in the Probe, which are not present in the Irrelevant, and, also, not in the Target. Because classic ERP analysis methods, like peak-to-peak or base-to-peak, would not fully reflect or measure the Probe’s multi-cycle oscillations, time frequency analyses (ERSP and ITC) were used, as outlined below.

To analyse EEG data in the time-frequency domain, the following two transforms were used: Event Related Spectral Perturbations (ERSP) and Inter-Trial Coherence (ITC). These were calculated, using a fast Fourier transform, with a baseline correction at -100ms to 0ms. ERSP calculates the average changes, relative to baseline, in power at each frequency at each time point, across all trials (Delorme & Makeig, 2004). ITC measures phase consistency between trials, determining the extent to which trials are phase-locked, at each time point and frequency (Makeig, Debener, Onton, & Delorme, 2004).

#### Time Frequency Window Placement

Time frequency analyses were measured, over two time windows, using orthogonal contrast time window placement (Kilner, 2013); (Kriegeskorte, Simmons, Bellgowan, & Baker, 2009); (Brooks, Zoumpoulaki, & Bowman, 2017); (Bowman, Brooks, Hajilou, Zoumpoulaki, & Litvak, 2020). In a similar way to our ERP analysis of the time domain, the time window for the Region of Interest that we used to measure ERSP and ITC, was identified based on aggregated power and coherence.

For group-level Time Frequency analysis, the placement of the critical time window (i.e. the highest 100ms interval in the broader time window of 0ms to 1200ms) for measuring ERSP/ITC was calculated using the average of power/coherence of all single trials of all participants (i.e. the aggregated grand average of all trials, across all participants) from both Probe and Irrelevant conditions. For participant-level Time Frequency analysis, the placement of the critical time window (i.e. the highest 100ms interval in the broader time window of 0ms to 1200ms) for measuring ERSP/ITC was calculated using each individual participant’s average of power/coherence (i.e. the aggregated ERP of all trials, for a single participant) from both Probe and Irrelevant conditions. Thus, both methods for time window placement were calculated independently of the contrast that is statistically tested.

Next, these orthogonally derived time windows could be employed to measure ERSP and ITC separately, in the Probe and Irrelevant conditions. In keeping with previous studies (Delorme & Makeig, 2004), the EEGLAB time-frequency function *newtimef* was used to calculate the ERSP and ITC for each condition. Each condition (i.e. Probe and Irrelevant) would supply this function with a matrix that contains its respective time-points and trials. The *newtimef* function would process these two input matrices and calculate two output matrices, which represent the difference in the power and coherence; the first output matrix comprised the difference in power (i.e. ERSP) between Probe and Irrelevant conditions, and the second comprised the difference in coherence (i.e. ITC) between Probe and Irrelevant conditions.

#### Time Frequency Statistical Inference

We summed across all values that were greater than zero, in the (orthogonally-derived) time window for the frequency range of interest, to give us a measure for each of the ERSP and ITC transforms for each condition. For this summation process we derived two difference (Probe minus Irrelevant) measures – one for power (i.e. ERSP) and another for coherence (i.e. ITC) – which would become the True Observed Values. Just as we had done in the time domain, we used a randomisation (Monte Carlo permutation) procedure to generate two Null hypothesis distributions: one for power and one for ITC (as calculated by the summation process, outlined above, across the orthogonally derived time windows). For each participant, we calculated *p*-values for both power and ITC transforms.

#### Detrending

Drift in EEG data can be a significant problem for the selection of an analysis window based upon identifying an interval with an extreme mean amplitude. For example, if the signal is drifting upwards, it is likely that the most positive-going window is at the end of the time series, and this selection would be made because of the drift rather than the amplitude of any particular component in the time series. Thus, in order to successfully use the aERPt or AGAT approaches, one needs a means of mitigating drift. The standard method for dealing with drift in EEG data would be to employ increasing high-pass filtering. However, filtering strategies may introduce temporal distortions in the signal, especially for low frequency P3 components that we are studying, where the amplitude of the P3 starts to reduce as the frequency cut-off increases. Therefore, we sought a method for mitigating drift, without introducing temporal artefacts. This was achieved through a detrending technique, whereby a linear trend line was fitted to the combined probe/irrelevant average and then this line was subtracted from every trial of both conditions.

Finally, baseline correction was performed after detrending, otherwise, we could be artificially lowering the late-components and increasing the early-ones (i.e. tilting the data, end-down and start-up).

## RESULTS

### Target Question

As previously explained, at the end of each RSVP stream, the participant was required to answer two questions. The first question related to the finishing-item and the second related to Target-recognition. Out of 75 times that each participant was randomly presented with the Target face, the average Hit rate for the group was 72.6% (*M* = 54.43, *SD* = 17.87), and out of the remaining 150 other trials in which the Target was not presented, the False-Positive rate was 3.3% (*M* = 4.93, *SD* = 4.12). The resulting discrimination measure (*d’* = 2.494) suggested that participants performed the Target task well, and no participants were excluded due to low discrimination.

### Probe/Irrelevant Questions

As previously discussed, at experiment end, the participant was given a recognition memory test, to determine if the three Probes and/or the three Irrelevants were perceived/recognised. This memory test consisted of 12 questions, appearing randomly, where each question accompanied an image that may or may not have been included in the experiment’s three blocks. Six questions related to the presence of the paired Probe and Irrelevant Lecturer faces that were included in the three blocks (i.e. one pair of Probe/Irrelevant, per block), and the other six questions related to random Probe and Irrelevant (Lecturer) faces that were not included in the experiment.

As before, the response to each of these 12 recognition/memory tests were handled in two parts: firstly, what is the participant’s confidence rating of how often each face was presented (i.e. the Probe/Irrelevant that were present, and the Probe/Irrelevant that were absent), and secondly, a confidence rating of how well the participant knew each of the 12 faces, prior to the experiment. The responses to both of these confidence ratings used a scale of 1 to 5, where 1 is “Never”, 2 is “Once or twice”, 3 is “Few times”, 4 is “Many times” and 5 is “A lot”. Note, for the purposes of statistical comparison, 1 out of 5 (i.e. Never) is equivalent to 0% and 5 out of 5 (i.e. A lot) is equivalent to 100%. Thus, 2 out of 5 = 25%, 3 out of 5 = 50% and 4 out of 5 = 75% (see Appendix B.1 of (Hajilou, 2020) for the full results).

### Overall Probe/Irrelevant recognition

The three Probe (familiar-lecturer) faces that were included in the experiment were reported to have been seen 33.9% of the scale (Mean confidence rating of 2.4 out of 5), and participants reported a very high (pre-experimental) familiarity of 94.6% with these Lecturer faces (4.8 out of 5). When comparing this to the (absent) Probe faces that were not included in the experiment, participants reported a similar high (pre-experimental) familiarity of 79.2% (4.2 out of 5), and only reported seeing these ‘absent’ familiar-lecturers 3.6% of the scale (1.1 out of 5), which is approximately one-ninth of the Lecturers that were, indeed, included in the experiment.

The three Irrelevants (*unknown*-lecturer) faces that were included in the experiment were reported to have been seen 4.8% of the scale (1.2 out of 5), and, similarly, the absent Irrelevant faces that were not included in the experiment were reported to have been seen 2.4% of the scale (1.1 out of 5). Finally, participants reported an imperceptible (pre-experimental) familiarity of 0% with all the Irrelevant/*unknown*-lecturer faces (1.0 out of 5).

As we were comparing Probe faces with Irrelevant faces, it was encouraging to discover that Probes were reported to have been seen 33.9% of the scale (*M* = 2.4; *SD* = 1.2504), which was seven times more than Irrelevants that were reported 4.8% of the scale (*M* = 1.2; *SD* = 0.428). Note that both conditions (Probes and Irrelevants) were, in fact, presented an equal number of times. The mean confidence rating of the main comparison conditions, for all participants, reveals a highly significant difference between the Probe (known-lecturer) faces and the Irrelevant (unknown-lecturer) faces, using pair-wise comparison (*M* = 1.1714, *SD* = 1.5122), *t*(13) = 2.898, *p* = 0.0125, *d* = 1.2545).

### EEG Analyses

We were interested in the EEG data across all the midline electrodes (Pz, Cz and Fz), but in-line with (Kaufmann, Schulz, Grünzinger, & Kübler, 2011), we expect the strongest brain responses to familiar faces, to be recorded at the Pz electrode. Therefore, we will start by focusing on the Pz electrode, reporting Time and Frequency domain analyses (at group and participant level), before reporting the same analyses at Fz and Cz.

#### Pz Electrode

At group-level, the grand average ERPs of all three critical stimuli (i.e. Target, Irrelevant and Probe), at the Pz electrode, revealed a clear difference between conditions (see Figure 2). The Target was task-relevant, so it elicited a large classic P3. The Irrelevant (an unknown face), did not present a similar pattern to the Probe or Target. This was as expected, because non-salient stimuli were unlikely to breakthrough into conscious awareness, due to the high presentation rate of the RSVP streams. Although, the Irrelevant exhibits an interesting negative deflection, peaking at 300ms, which we discuss later. In a similar fashion to the famous faces experiment (Alsufyani, et al., 2019), the Probe (familiar face) exhibited evidence of a continuous oscillatory pattern, within a 280 to 620ms time frame (and observed frequency of approximately 3-4 Hz) and a late positivity.

**Figure 2.**
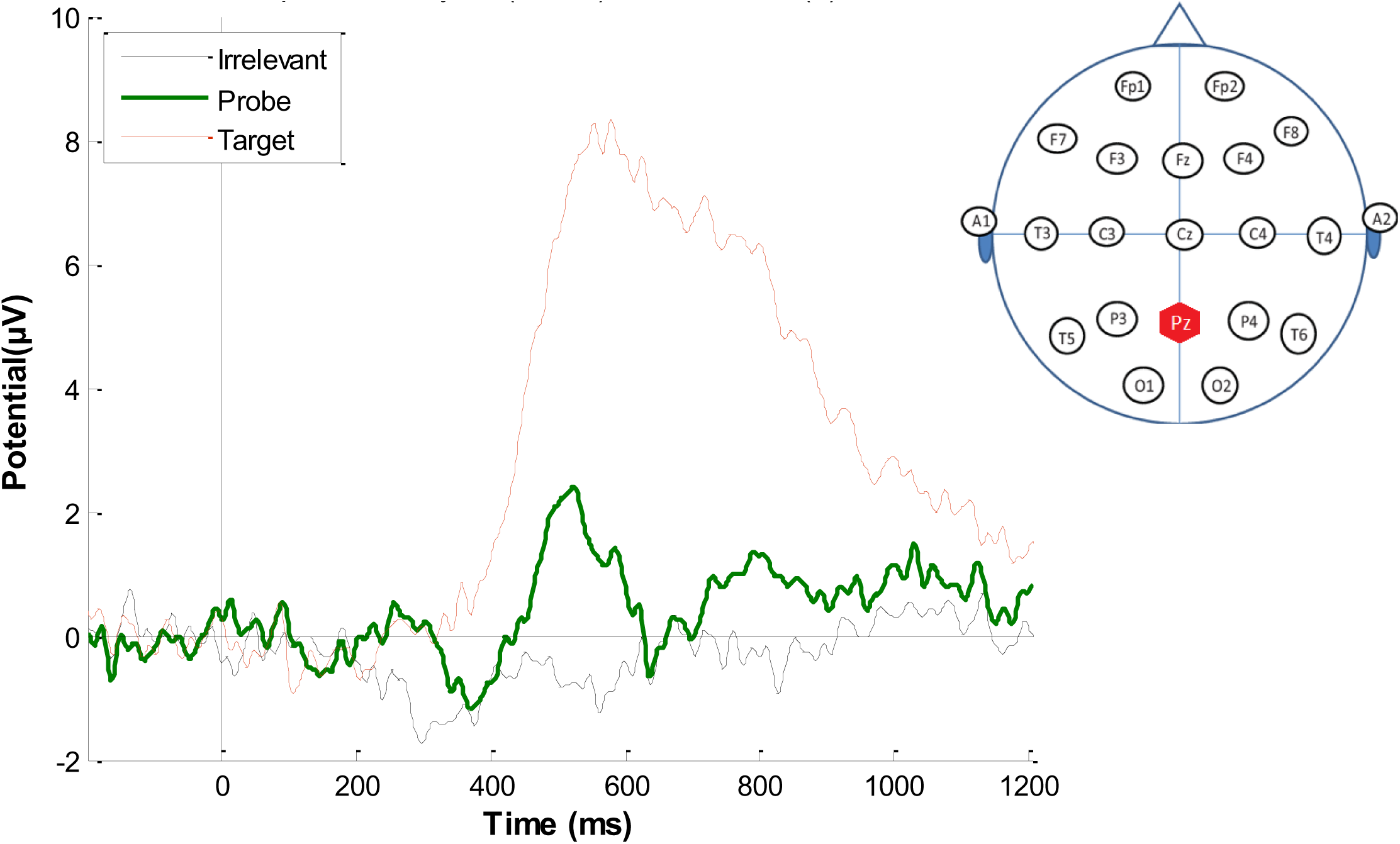
Grand average ERPs elicited by the three critical stimuli (Irrelevant, Probe and Target), at Pz, showing a classic P3 pattern for the **Target** (in red, peaking at +8.5µV), an oscillatory pattern for the **Probe** (in green, with an observed frequency of approx. 3-4 Hz) along with a late positivity, and a different pattern for the **Irrelevant** (in black, containing an interesting negative deflection, peaking at 300ms) that is distinct from the Probe and Target. As before, Target was the stimulus that the participant was instructed to look for, whereas, they were not informed of the presence of the Probe (familiar lecturer face). The oscillatory pattern for the Probe suggests a significant difference with the Irrelevants (unknown lecturer faces).

#### Group-level Analysis, at Pz

Within the a-priori P600f time-frame, the AGAT method independently identified an orthogonal contrast 100ms time window, at 453 to 553 (see Figure 3), and our statistical tests produced a highly significant difference between the Probe (*M* = 2.0086, *SD* = 2.0809) and Irrelevant (*M* = -0.4146, *SD* = 1.3911), at the Pz electrode (*M* = 2.4232, *SD* = 2.7311), *t*(13) = 3.3198, *p* = 0.0055, *d* = 1.3691.

**Figure 3.**
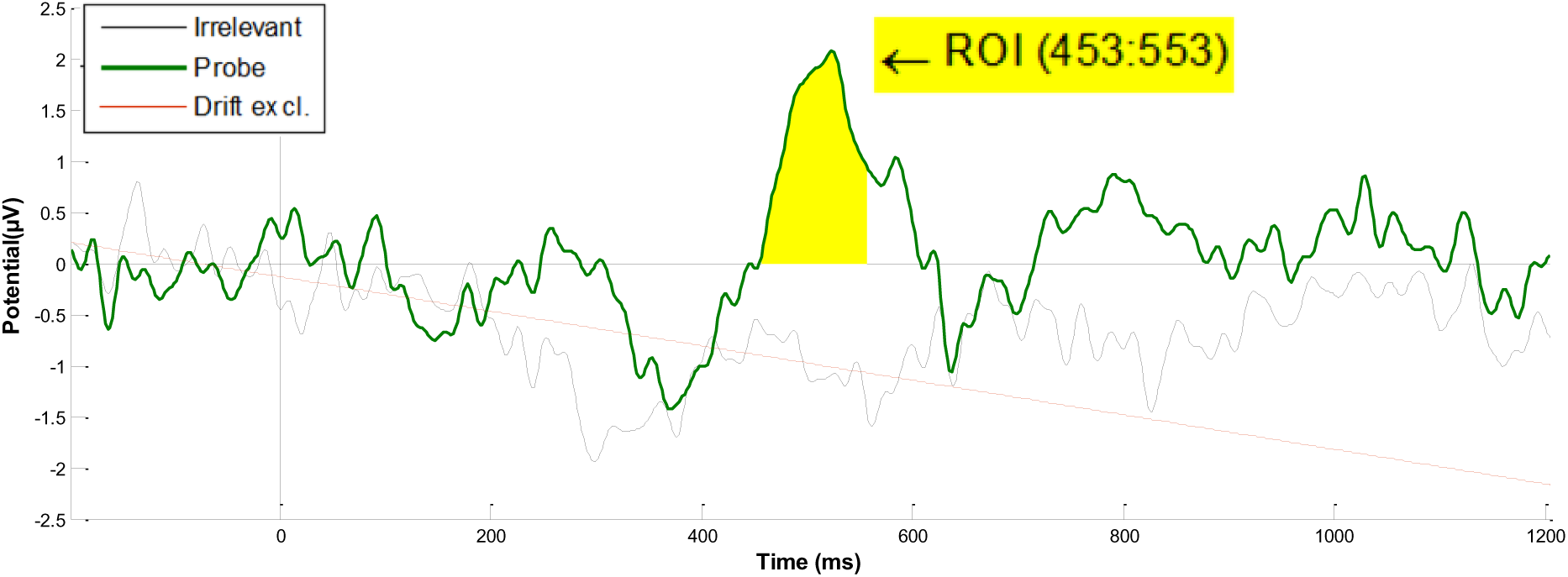
Grand average ERPs elicited by Probe and Irrelevant (i.e. the main comparison conditions) at **Pz**, showing an oscillatory pattern for the Probe condition (in green), which does not exist for the Irrelevant condition (in black). The linear Drift has been excluded (in red, at -9.7% to the vertical), with a detrending method. The window identified by the AGAT method is shown in yellow. Even though participants were not informed of the presence of the Probe (familiar lecturer face), statistical tests show a highly significant difference between Probe and Irrelevant, for **P600f** (t(13) = 3.3198, p = 0.0055, d = 1.3691). However, the same statistical test on N400f was not significant (t(13) = -1.9035, p = 0.0794, d = 0.4532).

Unlike the famous faces experiment (Alsufyani, et al., 2019), the same statistical tests on the N400f component did not result in a significant difference between the two conditions: (*M* = -0.57, *SD* = 1.1205), *t*(13) = -1.9035, *p* = 0.0794, *d* = 0.4532. Furthermore, participant-level statistical tests (i.e. Monte Carlo permutation) on the N400f component confirmed that none of the 14 participants (0%) showed critical-significance (0.05 alpha level) between Probe and Irrelevant (Mean *p*-value = 0.5744, *SD* = 0.2634). As can be seen in Figure 3, the unusual negativity in the Irrelevant condition (peaking at approximately 300ms, and overlapping the Probe condition’s N400f component) meant that the results of our statistical tests on N400f were not significant. In the remainder of the paper, we focus on the analysis of the P600f component.

#### Per-Participant Analysis

We performed statistical analyses of the ERP data, to determine whether the elicited response by the Probe (i.e. familiar lecturer face) was significantly different from that elicited by the Irrelevant (i.e. unknown lecturer face), on a per-participant basis. As shown in Table 1, per-individual statistical tests of Pz electrode’s P600f component, resulted in 8 of 14 participants (57.1%) achieving significance (at alpha level *p* < 0.05, shown in green) for the Probe – Irrelevant comparison.

**Table 1.** Per-participant analysis, at **Pz**, for the P600f component, showing the mean amplitude values of the Probe and Irrelevant conditions, from the same 100ms time window, which was independently found using the aERPt method. Significant participants are shown in green. Out of 14 participants, eight (57.1%) achieved critical-significance.

| Participant | Probe (*M*) | Irrelevant (*M*) | *p*-value |
| --- | --- | --- | --- |
| 1 | -0.8440 | -2.4415 | 0.1660 |
| 2 | 3.4664 | 0.1121 | 0.0360 |
| 3 | 0.7447 | 1.5237 | 0.6510 |
| 4 | 2.0746 | 1.7694 | 0.4350 |
| 5 | 0.5265 | -1.9373 | 0.0480 |
| 6 | 1.9794 | 0.7962 | 0.3280 |
| 7 | 4.2431 | -0.1308 | 0.0020 |
| 8 | -1.1190 | 0.5188 | 0.8580 |
| 9 | 1.4428 | -1.2087 | 0.0080 |
| 10 | 2.0194 | -1.3143 | 0.0270 |
| 11 | 4.6768 | -1.5233 | < 0.0001 |
| 12 | 0.1841 | 0.5206 | 0.6010 |
| 13 | 2.6896 | -0.1072 | 0.0340 |
| 14 | 6.0366 | -2.3814 | < 0.0001 |

As shown in Figure 4, the majority of participants’ Probes elicited a clear positive deflection, within the aERPt identified highest positive 100ms time window (highlighted in yellow), of the P600f search window (300 to 900ms, based on prior-precedent from literature). However, relative to the Irrelevant, the Probe for five participants (nos. 1, 3, 4, 6 and 12) failed to show a significant positivity for P600f, and one participant (no. 8) failed because the Probe ERP was consistently below the Irrelevant ERP.

**Figure 4.**
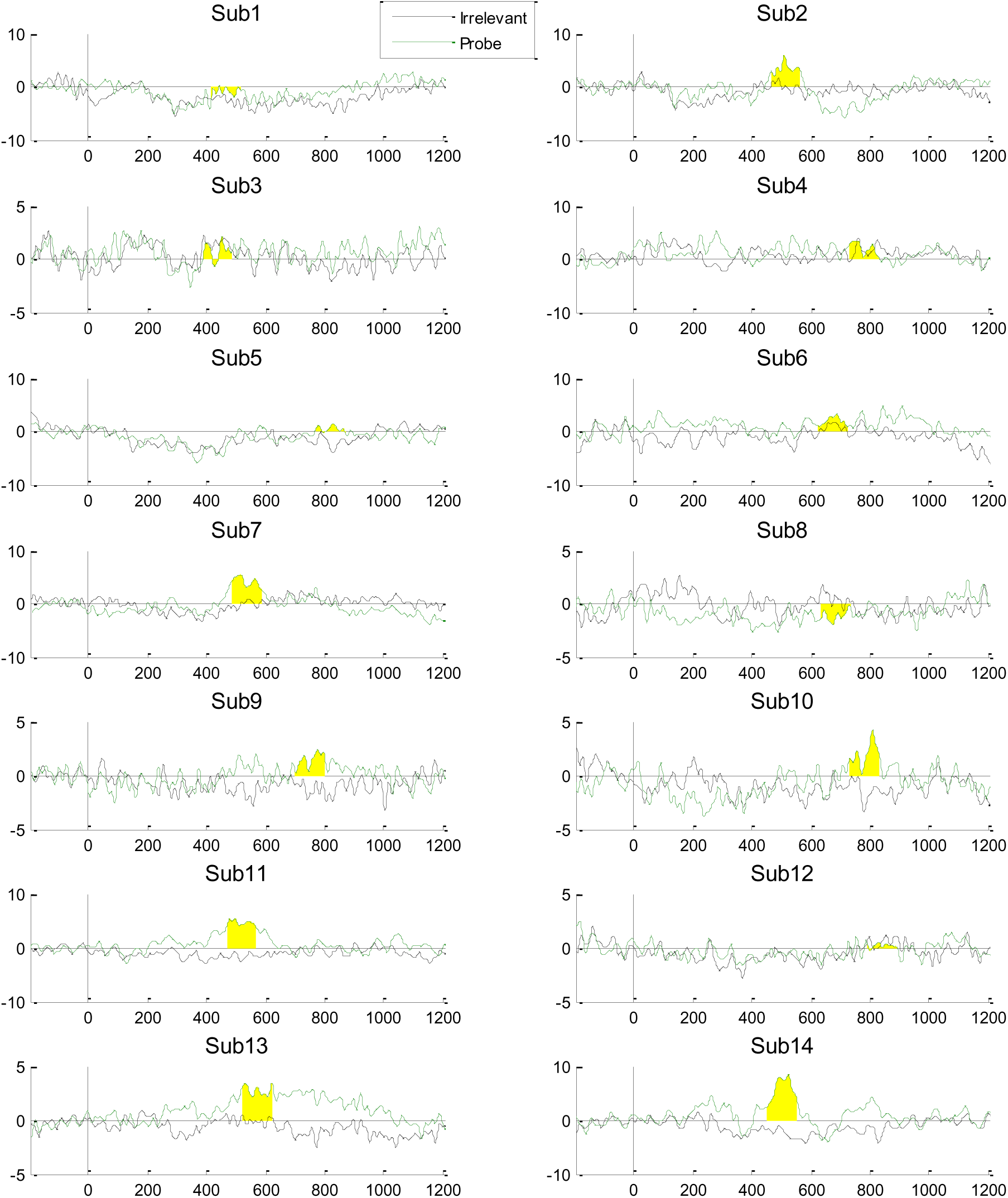
Per-participant Probe (in green) and Irrelevant (in black) ERPs, at the **Pz** electrode (x-axis represents Time in milliseconds, and y-axis represents Potential in microvolts). Each ERP shows the orthogonally identified highest positive 100ms time window (yellow highlight) for **P600f** (using the aERPt method), where 8 of 14 participants (57.1%) show a significant difference between the Probe and Irrelevant conditions.

#### Time Frequency Analysis (TFA)

As outlined in *Frequency Domain Analysis* (see Analysis section, above), to analyse the power and coherence of the EEG data, we have employed two Time Frequency transforms: Even-Related Spectral Perturbation (ERSP) and Inter-Trial Coherence (ITC), using EEGLAB’s toolbox (Delorme & Makeig, 2004). Whereas ERSP reflects the extent to which the signal power changes relative to the baselining window, ITC reflects the phase consistency (or synchronisation) between trials. ERSP/ITC changes in coherence enables us to quantify the multi-cycle oscillations that we had observed in the ERPs.

We applied a notch filter, between 7 and 9 Hz, in order to filter out any Steady State Visual Evoked Potential (SSVEP) set up by the RSVP stream. Thus, as long as there are no significant power increases at higher frequencies, we could fix the upper boundary of our analysis at 7 Hz (also noting that the lowest boundary is fixed by our standard high-pass filter, on 0.5 Hz). Even so, in addition to the fixed-boundary analysis window (0.5 to 7 Hz), we also performed the full ERSP/ITC analyses on the full frequency range (0.5 to 45 Hz), to assess the power/coherence changes at higher frequencies.

#### Group-level TFA

As seen in Figure 5, ERSP and ITC results of the AGAT of the Probe and Irrelevant conditions at Pz, exhibits a large power, respectively coherence, increase around 300 to 650ms (post-stimulus), mainly, at the low frequencies. To compare Probe and Irrelevant at the group-level, two measures were obtained for each participant, and a two-tailed paired *t*-test was used to perform statistical inference for ERSP and ITC. As can be seen in the grand-Probe versus grand-Irrelevant ERSP/ITC comparisons (see Figure 6), increases in power/coherence are predominantly evident in the grand-Probe condition, which suggests detection of the familiar lecturer face (ERSP > 4dB, and ITC > 0.4). However, the grand-Irrelevant condition lacks any significant power/coherence fluctuations, within the same time window, which implies little-to-no conscious or sub/liminal detection of the unknown lecturer face. Nearly all the power/coherence fluctuations are occurring in the lower frequency range (i.e. 0.5 to 7Hz).

**Figure 5.**
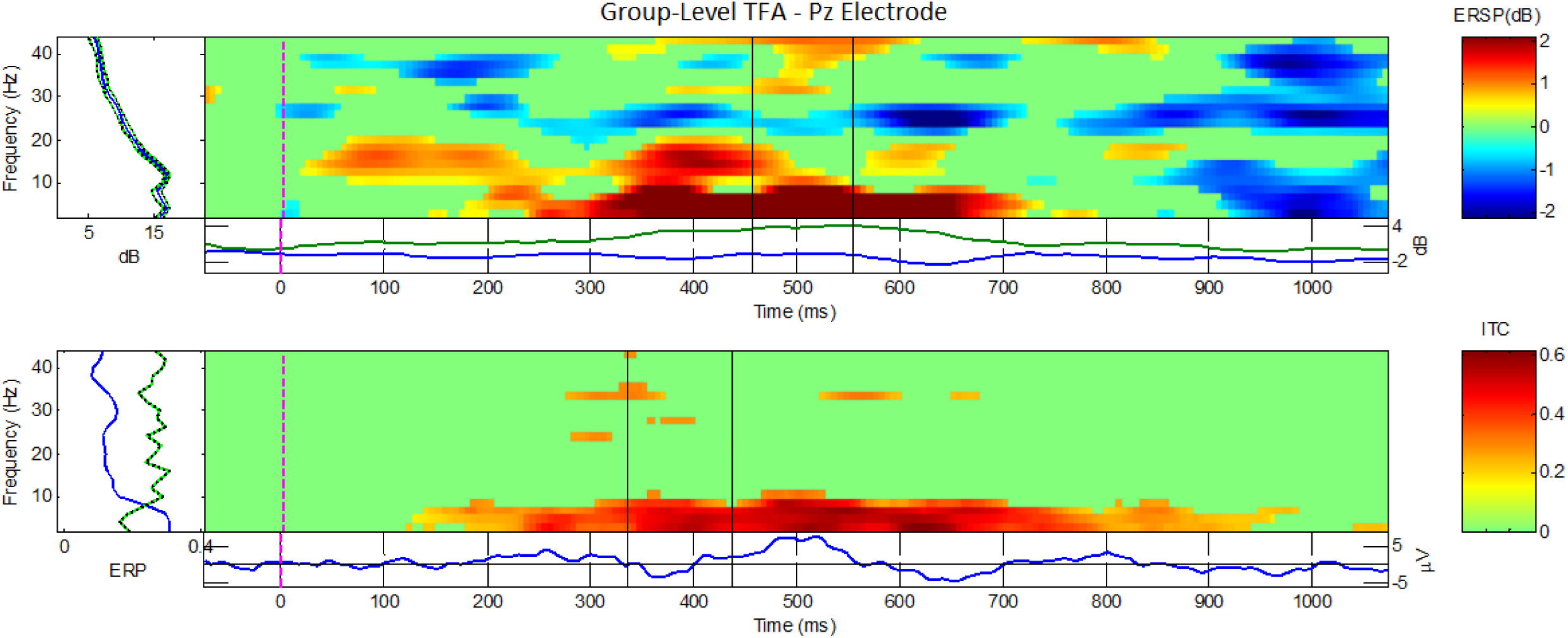
Group-level Time Frequency plots, at the Pz electrode, using the combined Probe and Irrelevant conditions, i.e. the AGAT. The top plot depicts the ERSP, and the bottom plot depicts the ITC. The independent window selection (using AGAT method) for ERSP produced a Region of Interest (ROI) at 461:561ms, and earlier ROI for ITC, at 334:434ms. Increases in power/coherence have been mostly concentrated in the 0.5 to 10 Hz frequency range, and are strongest in the ROI time frames (SSVEP has been filtered out, by applying a 7:9 Hz notch filter).

**Figure 6.**
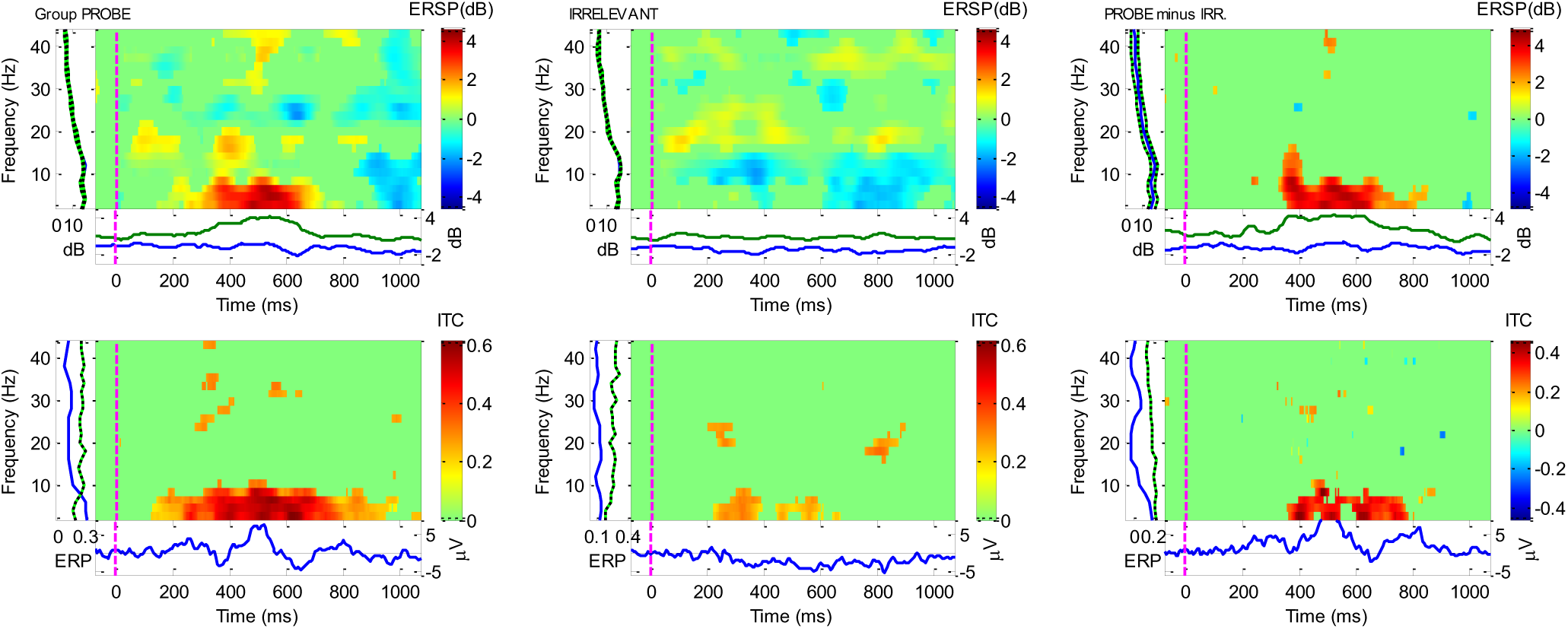
Group-level Time Frequency Analysis, at Pz electrode, for the difference between critical stimuli (Probe and Irrelevant), across the full frequency range (**0.5 to 45 Hz**), showing ERSP (top row) and ITC (bottom row). The first column of ERSP/ITC plots show the power/coherence changes in the grand-Probe condition, and the second column shows the same for the grand-Irrelevant condition. The third column is the difference between grand-Probe and grand-Irrelevant (i.e. Probe minus Irrelevant), which confirms group-level increases in power/coherence for the grand-Probe only: ERSP (t(13) = 2.9737, p = 0.0108, d = 1.0417), and ITC (t(13) = 1.8064, p = 0.0941, d = 0.7997). Note that at each frequency and time point, increases in power/coherence are in red; decreases in blue, and green indicates no significant change.

Over the full frequency-range (i.e. 0.5 to 45 Hz), the group-level analysis at Pz for ERSP was significant, at the AGAT defined window 461 to 561ms (see Figure 5), confirming a difference between the Probe and Irrelevant conditions (*t*(13) = 2.9737, *p* = 0.0108, *d* = 1.0417). For the group-level ITC over the same/maximum frequency range, our statistical test was not significant, at the AGAT defined window 334 to 434ms: (*t*(13) = 1.8064, *p* = 0.0941, *d* = 0.7997).

However, focusing on the narrower frequency-band (i.e. 0.5 to 7 Hz), the group-level analysis at the Pz electrode for ERSP was highly significant, at the AGAT defined window 484 to 584ms (see Figure 7), confirming a difference between Probe and Irrelevant (*t*(13) = 3.4769, *p* = 0.0041, *d* = 1.2649). For group-level ITC over the same (narrower) frequency range, our statistical test was also significant at the AGAT defined window 588 to 688 (*t*(13) = 2.322, *p* = 0.0371, *d* = 0.7442).

**Figure 7.**
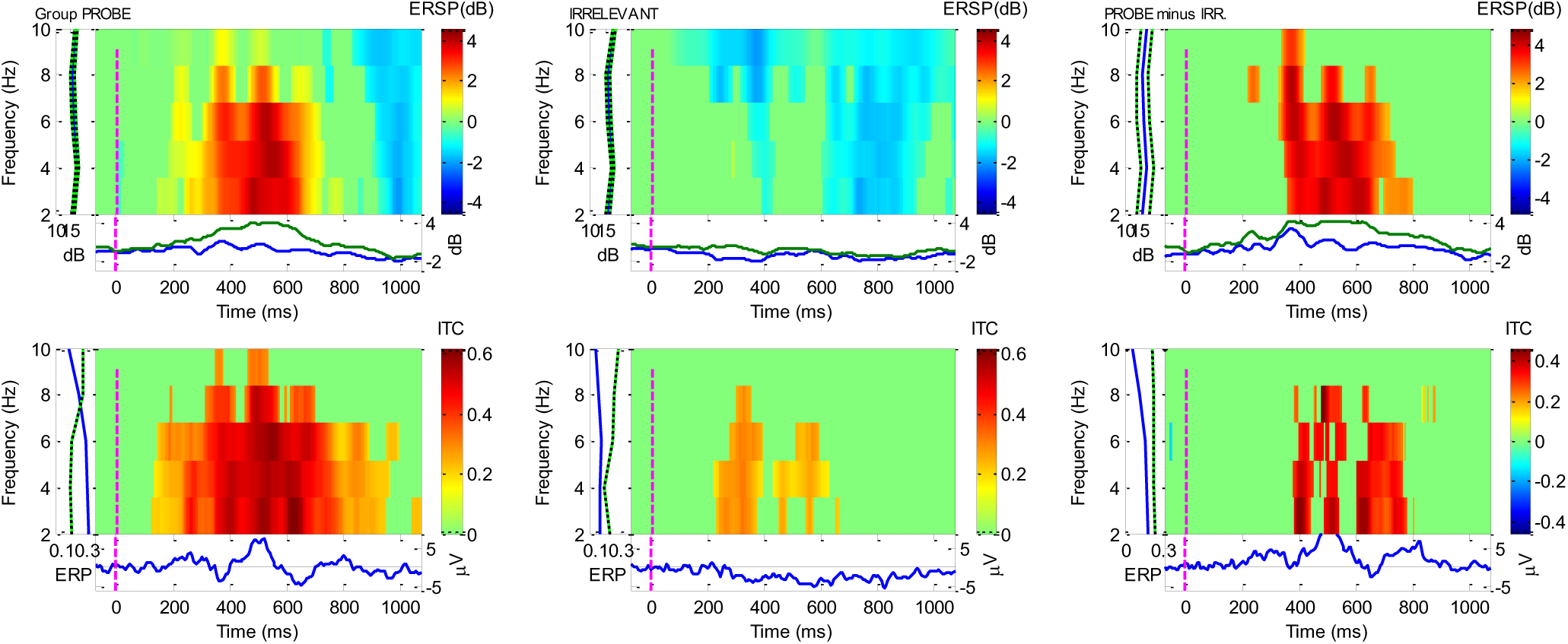
Group-level Time Frequency Analysis, at Pz, for the difference between critical stimuli (Probe and Irrelevant), at the narrower frequency band (**0.5 to 7 Hz**), showing ERSP (top row) and ITC (bottom row). The first column of ERSP/ITC plots show the power/coherence changes in the grand-Probe, and the second column shows the same for the grand-Irrelevant. The third column is the difference between grand-Probe and grand-Irrelevant (i.e. Probe minus Irrelevant), which confirms group-level increases in power/coherence for the grand-Probe only: ERSP (t(13) = 3.4769, p = 0.0041, d = 1.2649), and ITC (t(13) = 2.322, p = 0.0371, d = 0.7442). At each frequency and time point, increases in power/coherence are in red; decreases in blue, and green indicates no significant change.

#### Per-Participant-level TFA

Per-participant statistical inference – in the form of a randomisation test – confirmed the substantial increase in the Probe’s power (ERSP) and coherence (ITC), as compared to the Irrelevant. Statistical tests of the *narrower* frequency band (0.5 to 7 Hz), resulted in two orthogonal contrast time windows for each participant (ERSP average window: 496 to 569, and ITC average window: 413 to 513) and *p*-values that revealed a difference between the Probe and Irrelevant conditions (see Table 2). For ERSP, 10 out of 14 participants’ *p*-values (71.4%) were significant. Similarly, for ITC, 9 out of 14 participants’ *p*-values (64.3%) were significant.

**Table 2.** Per-participant Time Frequency analysis of power (ERSP) and coherence (ITC), at **Pz electrode**, using the narrower frequency range (**0.5 to 7 Hz**). For each participant, an orthogonal contrast time window was employed (using the aERPt method), and p-values were obtained for ERSP and ITC, by comparing the Probe and Irrelevant conditions, using a randomisation test. 10 of 14 ERSP p-values (71.4%) were significant, and 9 of 14 ITC p-values (64.3%) were significant.

| Participant | ERSP *p*-values | aERPt win. | ITC *p*-values | ITC win. |
| --- | --- | --- | --- | --- |
| 1 | 0.6733 | 703 | 0.0107 | 369 |
| 2 | <0.0001 | 508 | <0.0001 | 525 |
| 3 | 0.04 | 461 | 0.2493 | 162 |
| 4 | <0.0001 | 404 | 0.22 | 709 |
| 5 | 0.008 | 410 | 0.0147 | 520 |
| 6 | 0.0147 | 531 | 0.6213 | 260 |
| 7 | <0.0001 | 514 | 0.0293 | 428 |
| 8 | 0.2147 | 795 | 0.244 | 53 |
| 9 | 0.2907 | 473 | 0.0187 | 674 |
| 10 | 0.264 | 307 | 0.043 | 629 |
| 11 | <0.0001 | 490 | <0.0001 | 484 |
| 12 | 0.0253 | 559 | 0.284 | 53 |
| 13 | 0.0133 | 455 | 0.04 | 473 |
| 14 | <0.0001 | 334 | <0.0001 | 449 |

According to the above Frequency domain analysis (see Table 2), with the exception of participant 1, statistical test results of ITC appear to be closely correlated to the Time domain’s statistical tests of the ERP data (see Table 1), at per-participant-level, since both analyses produce significant *p*-values for participants 2, 5, 7, 9, 10, 11, 13 and 14.

Finally, even though we have justified fixing the upper boundary of our analysis at 7 Hz (i.e. due to SSVEP waveform, which required a notch-filter on 7 to 9 Hz), we confirmed that per-participant statistical analysis of the maximum frequency range (0.5 to 45 Hz), resulted in *p*-values that revealed a difference between the Probe and Irrelevant conditions (see Table 3). For ERSP, 7 out of 14 participants’ *p*-values (50%) were significant. As for ITC, 9 out of 14 participants’ *p*-values (64.3%) were significant.

**Table 3.**
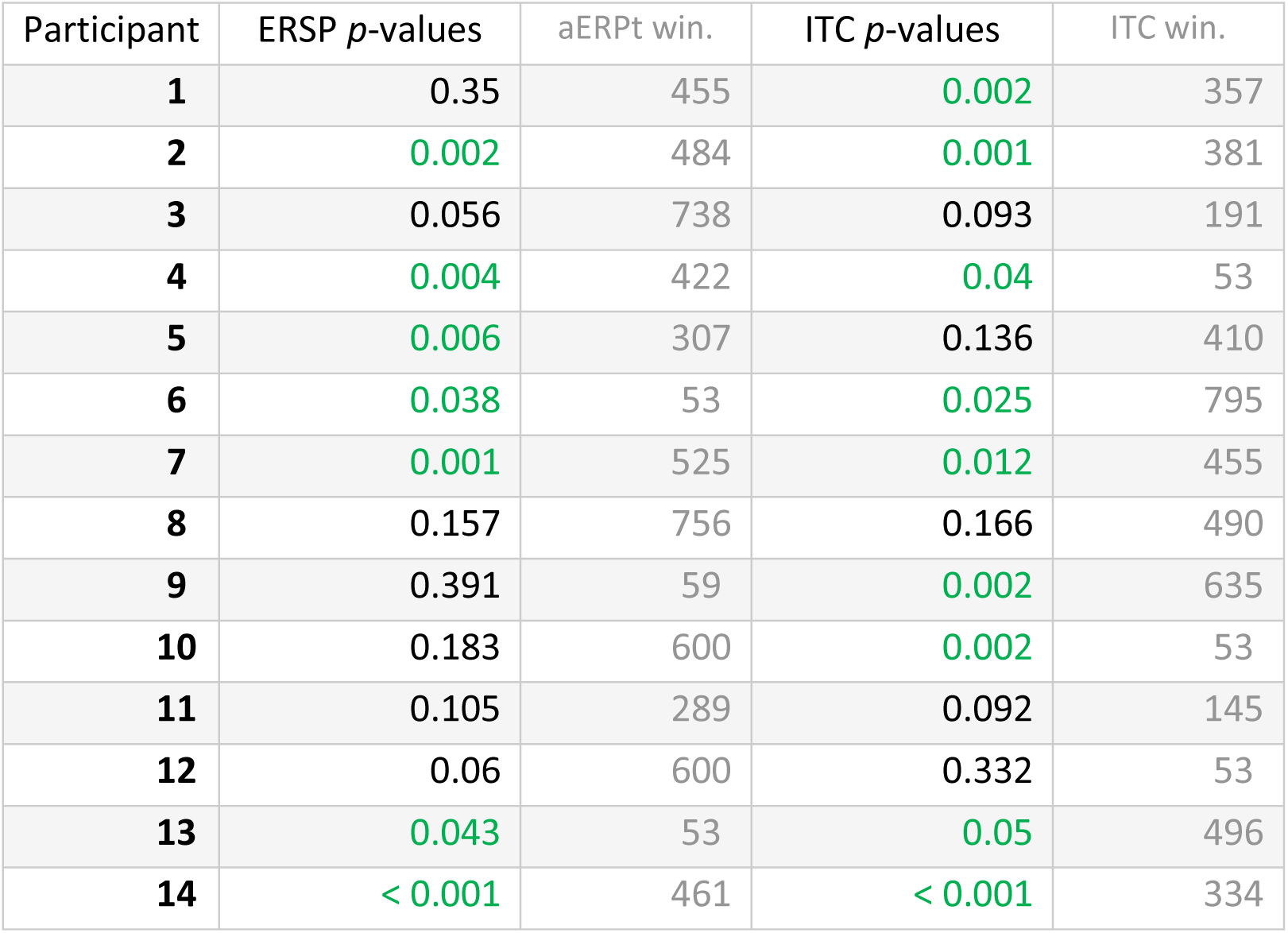
Per-participant Time Frequency analysis of power (ERSP) and coherence (ITC), at **Pz electrode**, using the maximal frequency range (**0.5 to 45 Hz**). For each participant, an orthogonal contrast time window was employed (using the aERPt method), and p-values were obtained for ERSP and ITC, by comparing the Probe and Irrelevant conditions, using a randomisation test. 7 of 14 ERSP p-values (50%) were significant, and 9 of 14 ITC p-values (64.3%) were significant.

| Participant | ERSP $p$-values | aERPt win. | ITC $p$-values | ITC win. |
| --- | --- | --- | --- | --- |
| 1 | 0.35 | 455 | 0.002 | 357 |
| 2 | 0.002 | 484 | 0.001 | 381 |
| 3 | 0.056 | 738 | 0.093 | 191 |
| 4 | 0.004 | 422 | 0.04 | 53 |
| 5 | 0.006 | 307 | 0.136 | 410 |
| 6 | 0.038 | 53 | 0.025 | 795 |
| 7 | 0.001 | 525 | 0.012 | 455 |
| 8 | 0.157 | 756 | 0.166 | 490 |
| 9 | 0.391 | 59 | 0.002 | 635 |
| 10 | 0.183 | 600 | 0.002 | 53 |
| 11 | 0.105 | 289 | 0.092 | 145 |
| 12 | 0.06 | 600 | 0.332 | 53 |
| 13 | 0.043 | 53 | 0.05 | 496 |
| 14 | < 0.001 | 461 | < 0.001 | 334 |

#### Other midline electrode sites

All the above Time and Frequency domain analyses focused on the Pz electrode, but we were also interested in the other two midline electrodes (Cz and Fz), to confirm that, in line with (Kaufmann, Schulz, Grünzinger, & Kübler, 2011), the strongest brain responses to familiar faces are recorded at Pz. The following analogous Time domain analyses of Fz and Cz, aim to find out if the P600f evoked by the Probe was significantly different from that evoked by the Irrelevant.

##### Fz electrode

At the group-level, the grand-average ERPs of the two critical stimuli (i.e. the Probe and Irrelevant), at the Fz electrode, revealed a clear difference between the conditions (see Figure 8). The Irrelevant did not present any feature/pattern similar to the Probe. Again, the Probe elicited a continuous oscillatory pattern, within a 200 to 620ms time frame. This waveform at Fz is similar to the oscillatory waveform at Pz (see Figure 3), and it confirms a large difference between the Probe (familiar lecturer face) and the Irrelevant (unknown lecturer face), at frontal midline electrodes. At this Fz electrode, an orthogonal contrast time window, for the highest positive (P600f) component, was found (using the AGAT method) at 434 to 534ms.

**Figure 8.**
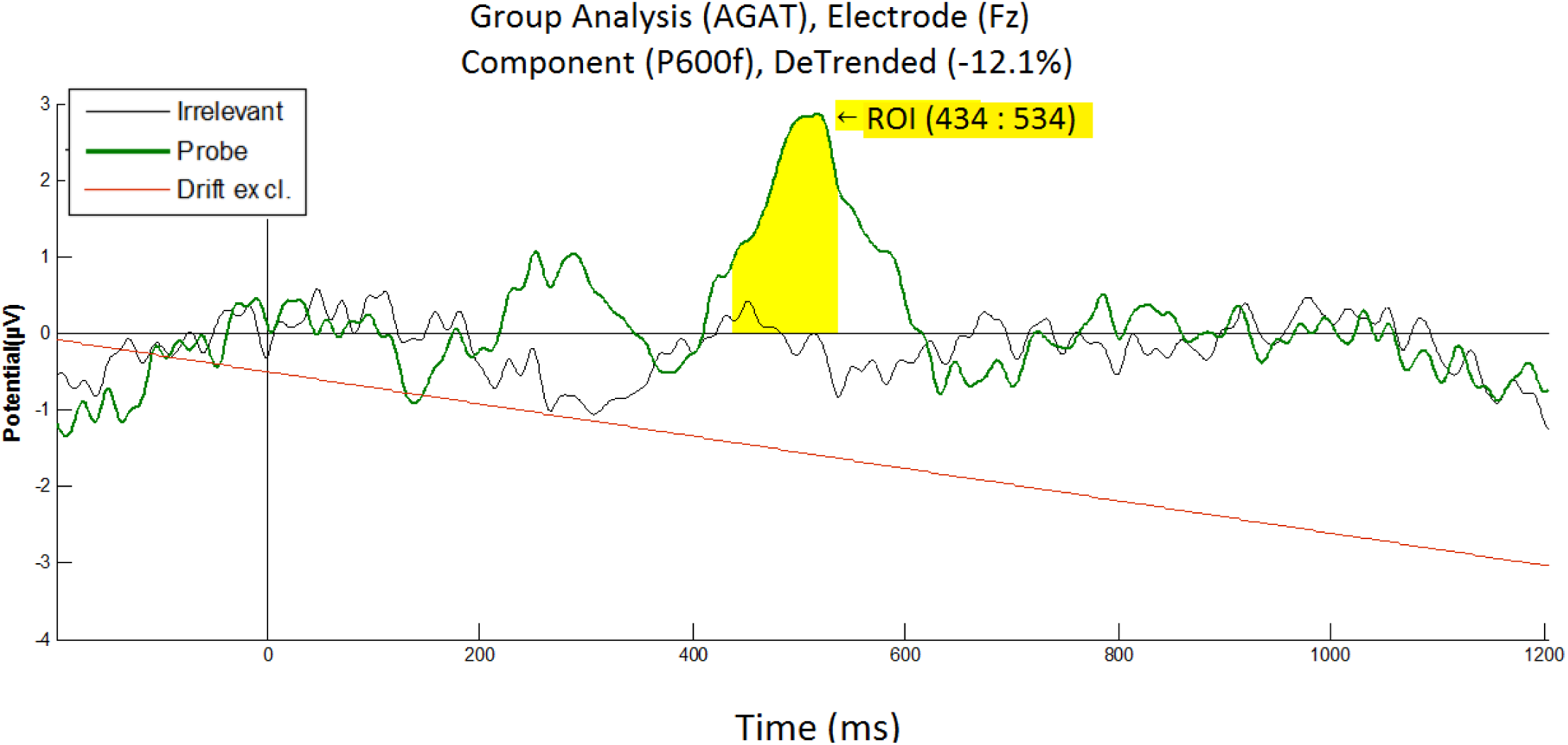
Grand average ERPs elicited by the Probe and Irrelevant, at **Fz**, showing an oscillatory pattern for the Probe (in green), which does not exist for the Irrelevant (in black). The linear Drift has been excluded (in red, at -12.1% to the vertical) with a detrending method. Baseline correction is between -100ms to zero. Even though participants were not informed of the presence of the Probe, statistical tests show a significant difference between Probe and Irrelevant for **P600f** (t(13) = 3.3151, p = 0.0056, d = 0.9834).

Statistical analyses at Fz – a paired *t*-test of the mean amplitudes of Probe and Irrelevant, across all participants – was run at the group level for the P600f component. Our statistical tests produced a significant difference between the Probe (*M* = 2.0696, *SD* = 2.4934) and Irrelevant (*M* = -0.0349, *SD* = 1.7156), at Fz: (*M* = 2.1045, *SD* = 2.3754), *t*(13) = 3.3151, *p* = 0.0056, *d* = 0.9834.

At the Fz electrode, per-participant statistical tests (i.e. Monte Carlo permutation) on the P600f component confirmed that 6 of 14 participants (42.9%) showed significant differences between Probe and Irrelevant (see Table 4). This confirms that results at Pz (i.e. 8 of 14; 57.1% – see Table 1), were better than at Fz, agreeing with studies (Kaufmann, Schulz, Grünzinger, & Kübler, 2011) that report stronger brain responses (to familiar faces) at Pz.

**Table 4.**
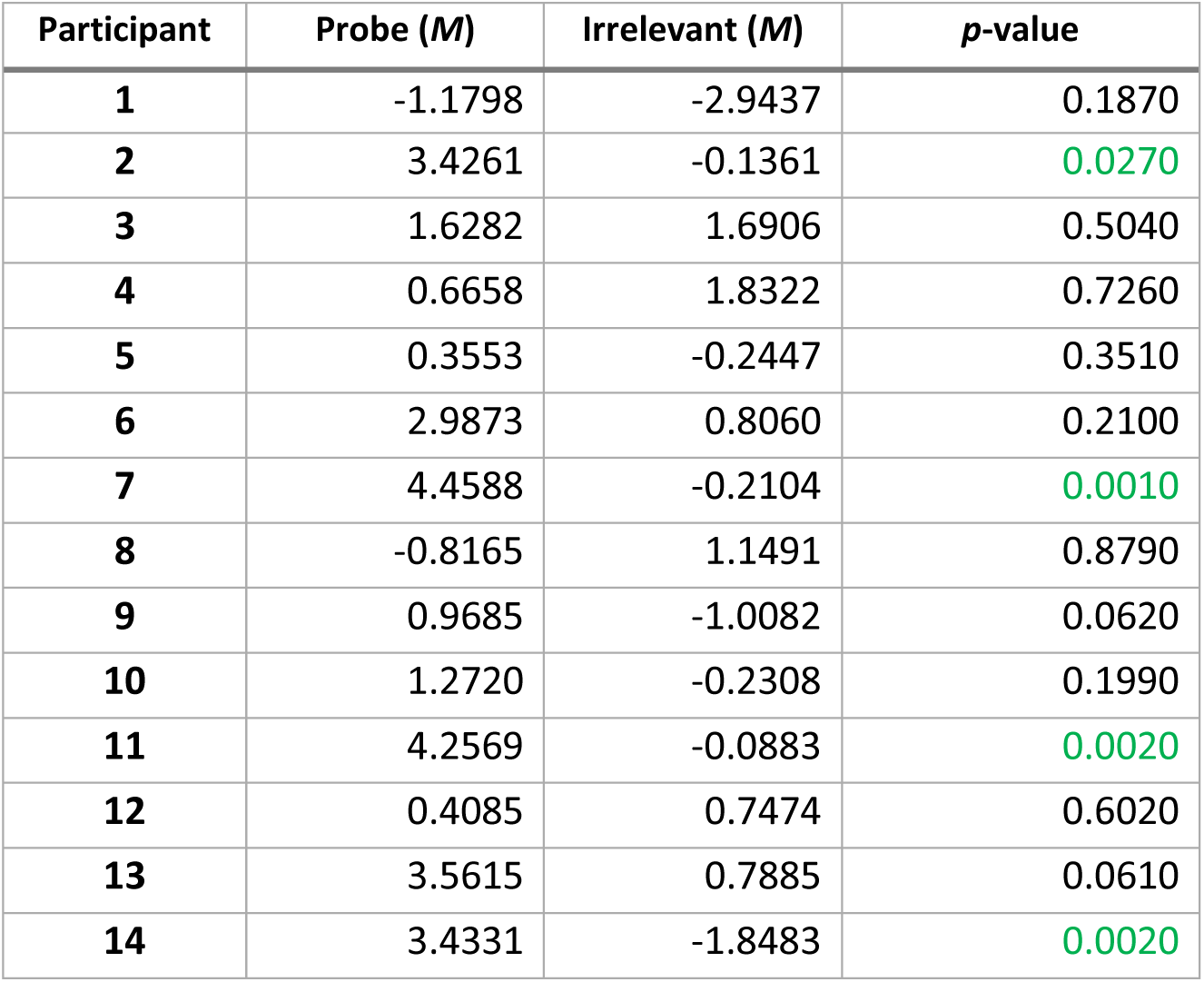
Per-participant analysis, at **Fz**, for the P600f component, showing the mean amplitude values of the Probe and Irrelevant conditions, from a 100ms time window, found using the aERPt method. Statistical tests on P600f resulted in 6 of 14 participants (42.9%) being significant, which is not as high as equivalent results at Pz (i.e. 8 of 14: 57.1%), confirming that Fz failed to show a stronger brain response when compared to Pz.

##### Cz electrode

The same group-level analysis that was carried out at Fz, was performed at Cz, revealing a difference between the Probe and Irrelevant conditions (see Figure 9). Once again, the Probe elicited a continuous oscillatory pattern, within a 220 to 720ms time frame, which was not present for the Irrelevant. This waveform at Cz is very similar to the oscillatory waveforms at Pz and Fz (see Figures 3 and 8, respectively). At this Cz electrode, an orthogonal contrast time window, for the highest positive (P600f) component, was found (using the AGAT method) at 438 to 537ms. A paired *t*-test for the P600f produced a significant difference between the Probe (*M* = 1.2271, *SD* = 2.0023) and Irrelevant (*M* = -0.4156, *SD* = 1.3367), at Cz: (*M* = 1.6427, *SD* = 2.1683), *t*(13) = 2.8346, *p* = 0.0141, *d* = 0.96494.

**Figure 9.**
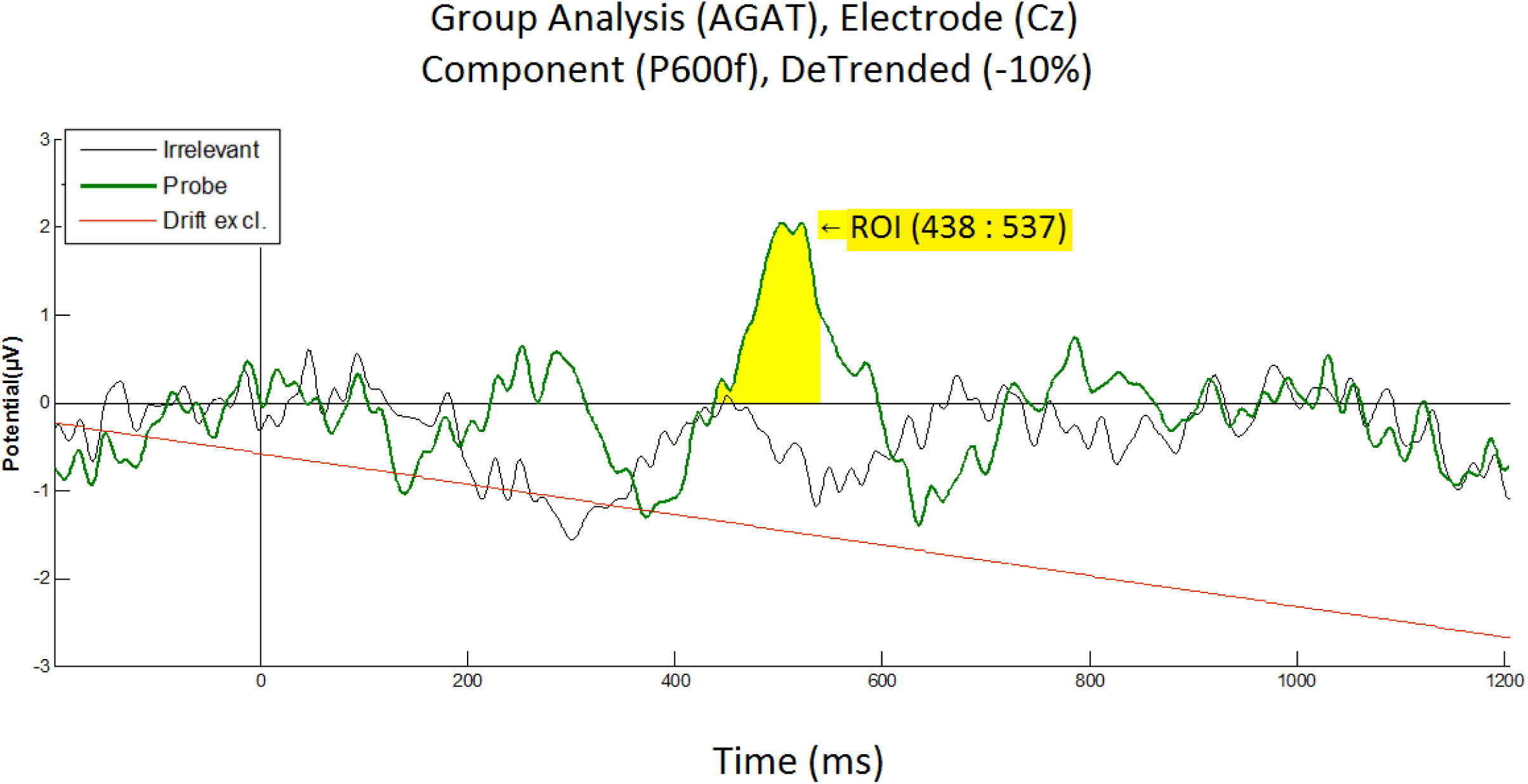
Grand average ERPs elicited by the Probe and Irrelevant, at the **Cz** electrode, showing an oscillatory pattern for the Probe condition (in green), for the Irrelevant condition (in black). The linear Drift has been excluded (in red, at -10% to the vertical) with a detrending method. Even though participants were not informed of the presence of the Probe (familiar lecturer face), a significant difference between Probe and Irrelevant, for **P600f** was observed (t(13) = 2.8346, p = 0.0141, d = 0.9649).

At the Cz electrode, per-participant statistical tests (i.e. Monte Carlo permutation) on the P600f component confirmed that 4 of 14 participants (28.6%) showed a significant difference between Probe and Irrelevant. Our per-participant results (see Table 5), confirmed that the effect was stronger at Pz (i.e. 8 of 14; 57.1% – see Table 1) than at Cz; once again, agreeing with studies that report stronger brain responses (to familiar faces) at the Pz electrode.

**Table 5.**
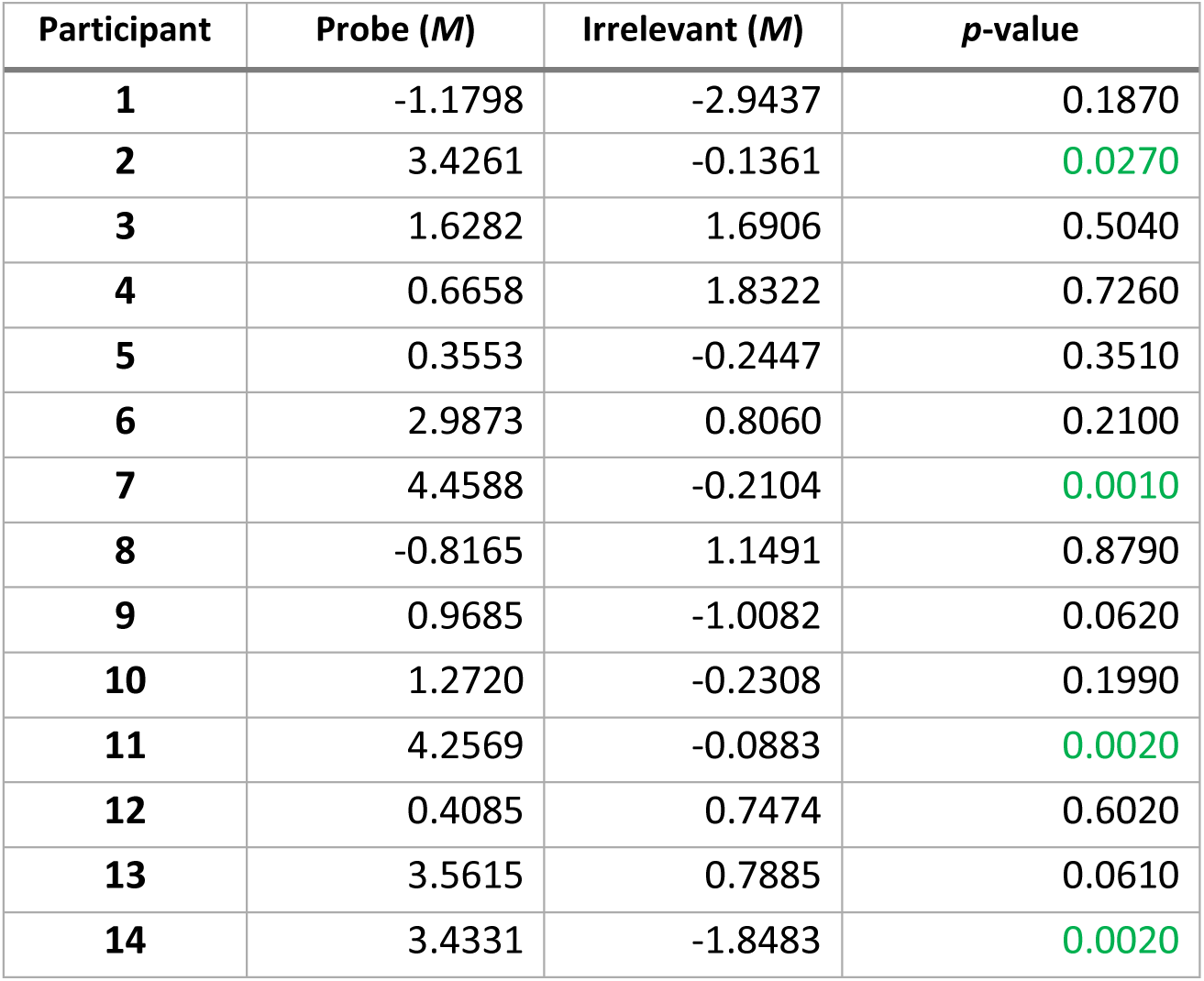
per-participant analysis, at **Cz** electrode, for the P600f component, showing the mean amplitude values of the Probe and Irrelevant, from the same 100ms time window, found using the aERPt method. Statistical tests on P600f resulted in 4 of 14 participants (28.6%) being significant, which is not as high as equivalent results at Pz (i.e. 8 of 14: 57.1%).

Finally, we have demonstrated that all three midline electrodes (Pz, Fz and Cz) have exhibited similar oscillatory waveforms, and that statistical tests showed significant differences between Probe and Irrelevant. Although our choice to focus on the Pz electrode was a priori (in-line with (Kaufmann, Schulz, Grünzinger, & Kübler, 2011)), we found that the strongest brain responses to familiar lecturer faces, was indeed recorded at the Pz electrode, as we observed in (Alsufyani, et al., 2019).

## DISCUSSION

The primary aim of the experiment presented here was to investigate whether faces that are personally known to an individual (familiar lecturer faces) can breakthrough into conscious awareness, and that event can be detected by EEG, on a group and per-participant basis, as successfully as our previous famous faces experiment (Alsufyani, et al., 2019). To do this, we performed statistical inference on ERP data (in the Time domain) and single-trial data (in the Frequency domain), to determine whether the evoked response by the Probe (familiar lecturer) faces were significantly different from that evoked by the Irrelevant (unknown lecturer) faces.

The key comparison was between Probe faces and Irrelevant faces, for which a significant difference was observed between the ERPs, at all three mid-line electrodes (Pz, Fz and Cz); see (Hajilou, 2020) for more details. However, unlike the prominent negative-and-positive deflections for the Probe (i.e. N400f, followed by P600f) we previously observed for famous faces (Alsufyani, et al., 2019), this experiment’s negative deflection (N400f) was muted, but its positivity (P600f) was equivalent (over a similar time frame of 300ms to 600ms). Additionally, even though there was again evidence of an oscillatory pattern, this second experiment’s ERPs are somewhat different and are typically weaker than those of the earlier famous faces experiment.

We closely mirrored the famous faces experiment’s design and analysis, with the main change being to the critical stimuli (i.e. replacement of the *celebrity* faces with *lecturer* faces). Even though the first (celebrity faces) experiment (Alsufyani, et al., 2019) established the viability of using faces to infer recognition in the RSVP paradigm, the next logical step was to investigate whether faces that are personally known to an individual would have a similar effect. After all, deception detection tests would seek to demonstrate a person under investigation’s recognition of somebody he/she has a personal relationship/familiarity with, rather than the non-partisan knowledge of a famous person.

In keeping with the famous faces experiment, participants were not informed that familiar faces may appear in the RSVP streams, and yet, our statistical tests confirmed the breakthrough events. Having been instructed to only look for the Target face, the inclusion of Probe faces (i.e. University lecturers) assessed participants’ ability to perceive incidental intrinsically salient faces. Statistically comparing brain responses to Probes and Irrelevants, in the time, as well as, frequency domains, enabled us to confirm differential perception of the Probe over the Irrelevants (i.e. unknown faces), at group and participant levels.

### Time Domain

At the Pz electrode, group-level analysis of ERPs confirmed the significance of the difference between the grand-Probe and grand-Irrelevant (*p* = 0.0055), and per-participant statistical analyses of ERPs confirmed that a total of 8 of 14 participants (57.1%) had significant *p*-values, revealing a difference between Probe and Irrelevant. Again, we confirmed that the brain response to the Probe at the per-participant level (8 of 14), achieved more participants at the Pz electrode, than Fz (*p* = 0.0056, and 6 of 14) or Cz (*p* = 0.0141, and 4 of 14).

### Frequency Domain

At the Pz electrode, group-level Time Frequency analysis, across the narrower frequency band (0.5 to 7 Hz), confirmed the significance of the difference between the grand-Probe and grand-Irrelevant for ERSP (*p* < 0.0041) and for ITC (*p* = 0.0371). Per-participant statistical inference, across the same frequency band (0.5 to 7 Hz), confirmed that 10 out of 14 participants’ *p*-values (71.4%) were significant. As for ITC, per-participant statistical inference showed that 9 out of 14 participants’ *p*-values (64.3%) were significant, confirming the difference between the Probe and Irrelevant conditions.

Thus, our experimental findings show substantial differences between the Probe and Irrelevant, which we believe result from the former stimuli reaching conscious awareness and generating pronounced electrical responses (as seen in the Probe ERPs), as suggested by our recognition memory experiments. However, there was an interesting new electrical response: a negative deflection to Irrelevant, peaking at 300ms, which we did not observe in the famous faces experiment (see for example, black line in figure 2).

It is noteworthy that participants did not report seeing the Irrelevant (unknown lecturer) faces, i.e. their recognition memory performance for Irrelevants was very poor. Accordingly, we have formed two theories to potentially explain this phenomenon. The first is that this posterior negativity may be related to subliminal registering (i.e. a covert response) of a repetition by the brain, and the second is that it may relate to an incongruity, between the Irrelevant and filler/distractor images, which was not present in the previous (famous faces) experiment, or indeed, both. More specifically, the Irrelevant images in the previous (famous faces) experiment were chosen randomly from the Distractor database, but the Irrelevant images in this (lecturer faces) experiment did not come from the Distractor database – they were (unknown) lecturer faces from a different University. In particular, to ensure a fair comparison with Probes, we used faces of academics that were unfamiliar to the participants. This necessarily, meant a “category” change between Distractors and Irrelevants. The key aspect of this change may just relate to changes in low-level visual features, since the Irrelevants were photos taken by the experimenter.

It also seems likely that this difference did not generate a breakthrough event for Irrelevants, since, as previously discussed, end of experiment, memory performance for Irrelevants was low. Finally and importantly, while the seemingly subliminal nature of this evoked response to the Irrelevants means it could be of scientific interest, the presence of such a response does not confound the logic of our experiment, although it may weaken the differences between Probe and Irrelevant that we observe.

### Size of effect

The per-participant statistical results were not as strong as those for the famous faces experiment. This is likely to be because the familiar (lecturer) faces do not have the level of salience of the famous faces. However, it may also be impacted by the previous exposure to the *images* of the famous *celebrity* faces (i.e. published photographs, which were frequently in the public eye, and surely seen by many participants, on many occasions and over a long period of time). In contrast, the *lecturer* faces of the experiment presented here, whilst having some personal salience to the participants, their *photographs* were seen for the first time, in the fast moving RSVP stream of images. In fact, the behavioural/recognition tests for both experiments support this premise, as participants in the previous famous faces experiment reported seeing the (celebrity) Probes with an average confidence rating of 3.4 out of 5 (60%), whereas, participants in the current experiment reported seeing the (lecturer) Probes with an average confidence rating of only 2.4 out of 5 (33.9%).

One possible explanation for weaker findings is that participants did not “see” all three of the Probe faces presented in the experiment. This may be because they did not have a strong familiarity with all three of the selected Probe faces. Consequently, an interesting line for future work is to develop a statistically valid method to “close-in” during the experiment on the Probe face for which the participant has the most familiarity. Of course, simply selecting the largest brain response post hoc would inflate the type 1 error rate. As a result, some consideration of the appropriate statistical procedure, probably through ground-truth synthetic simulations, is required to take such a closing-in procedure forward.

In addition future studies could take one step closer to the real-world scenario of concealed information tests, whereby, the hidden nature of the Probe critical stimuli would be to some extent revealed to the participants, at the start of the experiment. Having demonstrated that the breakthrough of familiar faces can be detected even when participants were not expecting to see celebrity-or-lecturer faces – because the presence of the Probe was concealed and the Target was the only task-relevant objective – we note that one could inform participants that, in addition to the Target (which will remain task-relevant), they may see a familiar lecturer face, in the RSVP streams. Naturally, one must not inform participants which lecture faces may appear in the experiment (or show them any photographs), but one can place their minds in a state where they are expecting to see the class of stimuli that the Probes will satisfy. This arrangement is closer to the real-life application of a deception detection test, in which the perpetrator would be fully aware of the purpose of the experiment (i.e. to find out if s/he is familiar with an accomplice). This strategic priming of the participants is likely to increase the effectiveness of the method.

### Countermeasures and Practicality

As previously emphasized, a major benefit arising from the fast presentation rate of the Fringe-P3 method is the demonstrated robustness against key countermeasures (Bowman, Filetti, Alsufyani, Janssen, & Su, 2014). However, two other issues that might be considered a problem for the Fringe-P3 method are, 1) attending away and 2) individual-specific perceptual thresholds. We discuss these in turn.

1. *Attending away*: participants would affect the results of the method, if they avoid (or fail to observe) images that are presented in the middle of the screen. They could do this by shifting their attentional focus away from the stream, with covert shifts of attention potentially being particularly problematic, since they would not be detectable with eye tracking. However, if participants are attending to the stream, a Steady State Visually Evoked Potential (SSVEP) will be observable at electrodes over visual cortex and since one knows the frequency of the stream, this steady state response can be efficiently detected. Thus, absence of an SSVEP would indicate noncompliance. Additionally, one could use the Target image Hit/False Alarm rate, as a further indicator of attentional engagement.
2. *Individual-specific perceptual thresholds:* we expect a difference between the Probe (known face) and Irrelevant (unknown face), but in order to reduce opportunities for countermeasures, the stream’s presentation rate must be high enough to only permit the Probe images to breakthrough into conscious awareness. Indeed, Probes may not be perceived if the presentation speed is too high, and Irrelevants may breakthrough if the speed is too slow. However, there are considerable individual differences in perceptual thresholds, which suggest that the optimal presentation rate will vary across participants. Whilst we have adopted a fixed presentation rate (e.g. a-priori SOA of 133ms), future studies could improve the breakthrough difference between the Probe and Irrelevant, by running a pre-experiment training session using a ‘staircase’ procedure, to determine the optimal presentation rate for each participant.

### The Fringe-P3 Method

It is notable that the form of the evoked response that we observe for probe stimuli does vary according to the type of stimulus presented. In particular, for face stimuli, we observe an oscillatory pattern in the theta frequency range. This, for example, is evident in the green time series in figure 2, and is an even more pronounced feature in our famous faces experiment (Alsufyani, et al., 2019). The practical value of the Fringe-P3 method is not impacted by the presence or absence of an actual P3 in these evoked response patterns; indeed, as we have shown here and in (Alsufyani, et al., 2019), analyses that use this oscillatory feature are particularly effective. However, these heterogeneous evoked patterns might challenge the reference to P3 in the name we have attributed to this method (i.e. Fringe-P3). However, for the Probe (green in figure 2), if one filtered out high frequencies (which amongst other things will make the negative deflection just after 600ms less sharp), and then subtracted out a (negative going) wavelet, centred on the negative deflection just before 400ms (which our famous faces data (Alsufyani, et al., 2019) suggests is the central phenomenon of the wavelet), one might end up with something that looks like a (weak) P3, peaking around the same time as the Target (red in figure 2). Thus, although further work is certainly required, it may be that the oscillatory feature in our faces experiment is “sat on top of” a P3. This would justify the term Fringe-P3 method still being applicable.

This paper’s experiment extended our earlier work, which, together, demonstrated that both highly evocative faces (i.e. famous faces experiment) and personally familiar faces (i.e. this experiment’s *lecturers*) can be employed in RSVP-based Fringe-P3 studies, and that highly familiar faces can breakthrough into conscious awareness, on a per-individual basis.

The results raise the possibility that one could apply our findings to the differentiation of deceivers and non-deceivers, in the application of crime compatriots, whereby, a suspect’s familiarity with a criminal/terrorist can be established using faces.

## Footnotes

1 If the Irrelevant can be detected, the electrical response to it can be increased, potentially making its response statistically indistinguishable from that to the concealed knowledge (the Probe), confounding the deception detection.

